# Evaluation of welfare indicators for companion parrots: a Delphi consultation survey

**DOI:** 10.1101/2024.06.20.599871

**Authors:** Andrea Piseddu, Yvonne R. A. van Zeeland, Jean-Loup Rault

## Abstract

Parrots can experience several welfare challenges when kept as companions. Despite their popularity no science-based guidelines are available to assess parrot welfare. The aim of this Delphi study was to evaluate welfare indicators that owners could use to monitor parrot welfare. One hundred and twenty-two potential welfare indicators (behaviours, body measurements, husbandry and management conditions) were sourced from a systematic literature review and by consulting an avian medicine specialist. They were presented to participants with expertise on parrots in two rounds of online survey. We identified 73 welfare indicators that could be used by owners to monitor the welfare of all/most parrot species. Abnormal behaviours and management conditions that allow parrots to express their natural behaviours were ranked among the most important indicators. Participants concurred with scientific evidence about the impact of diet, species susceptibility to develop behavioural problems, early life, and pre-acquirement experiences on parrot welfare. When inquired about the suitability of species as companions, participants indicated seven small-sized species as most suitable to keep as a companion parrot, while cockatoos, critically endangered, and highly trafficked species were evaluated as those that should not be kept as companions. These findings could be useful to monitor and improve parrot welfare.

## Introduction

Parrots are popular companion animals, appreciated for their intelligence, beauty and vocal ability as well as for the emotional support that they provide to their owner^1–3^. However, parrots can face several welfare challenges when kept in captivity. Poor welfare can arise due to a lack of knowledge or neglect of parrots’ biological needs and can manifest through health issues (e.g. obesity, atherosclerosis, fungal and bacterial infections) and behavioural problems, including excessive screaming, aggression, self-injurious behaviours and stereotypies^4–7^. A lack of cognitive stimulation, opportunities to forage, social interactions, locomotor behaviour and the provision of unbalanced diets are considered the main risks to parrot welfare^7–9^. Moreover, inappropriate human-parrot interactions can cause companion parrots to become aggressive, fearful, and excessively vocal^5, 10^. The emergence of these behavioural problems can have negative consequences on the parrot-human relationship and is considered one of the main causes of relinquishment for companion parrots^5, 10, 11^. Data on relinquishment is difficult to find; however, according to the limited information available, a high number of parrots are relinquished every year due to the difficulty of keeping them and fulfilling their needs^12, 13^. Considering the increasing popularity of parrots as companion animals and their longevity^2, 4, 14, 15^, and the serious and multiple welfare challenges faced in relation to the complexity of their needs, evidence-based guidelines for assessing and improving parrot welfare are urgently needed. More specifically, it is necessary to identify scientifically valid welfare indicators that ideally could be used by owners routinely. This would benefit both parrots and owners in several ways: it would inform about the appropriateness of the husbandry and management conditions provided; it would allow to regularly assess the parrot’s welfare state; and as a prophylactic measure, it could prevent the emergence of illness or behavioural problems, thus enhancing the possibility to maintain a good parrot-owner relationship and reducing the risk of parrot relinquishment.

At current, a large amount of potentially valuable information regarding parrot welfare is based on expert knowledge or experience and often reported through books or magazine articles, whereas comparatively less information is derived from experimental scientific studies. Nevertheless, we found a number of potential parrot welfare indicators and risk factors in a recent systematic literature review^16^. However, this review revealed a high risk of bias in the peer-reviewed scientific studies gathered, making it difficult to ascertain both the internal and external validity of the findings and therefore requiring an alternative process of validation^16^.

For this purpose, the so-called Delphi method is considered a suitable solution. This method consists of consulting a panel of experts that provide their opinion on a determined topic through multiple rounds of survey ^17^ and is based on four features: 1) anonymity, to avoid the risk participants could influence each other; 2) iteration, to allow participants to re-assess their judgment through multiple rounds; 3) controlled feedback, to inform about the responses provided by other participants; and 4) statistical aggregation of the group response^18^. The Delphi technique is a well-established method to assess content-related validity^19^ by reaching consensus among participants, with the assumptions that a group of individuals with different types of expertise and who anonymously and independently provide their opinion is better in decision-making than a single individual^20^. The consensus obtained through this standardized process can be considered meaningful when group stability is achieved^21^, meaning that the results of successive rounds of survey should not statistically differ^22^. Delphi consultation surveys have been used for decision-making in several research fields including animal welfare science, for example to identify indicators of welfare in laboratory-housed macaques^23^, mice^24^, captive reptiles^25^, shelter dogs^26^, horses^27^, farm pigs, dairy cattle and laying hens^28^ and meat chickens^29^. To our knowledge, the Delphi technique or other validation techniques based on experts’ consultation have thus far not been used to study parrot welfare.

The objective of this study was to conduct a Delphi consultation survey in order to screen indicators related to parrot welfare previously identified in the systematic scientific literature review or based on expert knowledge. We aimed to identify which welfare indicators are considered by experts as 1) valid for most parrot species; 2) feasible to use in practice by caregivers; 3) the most important; and 4) which factors are considered to impair parrot welfare. In addition, considering the large variety of parrots species, we aimed to determine which species, according to experts, are best suited to be kept as companions and which should not.

## Material and Methods

### Ethical consideration

The project was assessed by the Ethics Committee of the Medical University of Vienna, which determined that, in accordance with the Good Scientific Practice guidelines and relevant national legislation, an ethical approval was not required for this study. All participants gave their informed consent before participating in both the first and second round of the survey. Only one researcher (AP) was able to trace back the identity of each participant and their responses, ensuring quasi-anonymity^30^. This was required due to the iterative nature of the Delphi method in order to create personalized survey for each participant in the second round based on their previous responses provided in the first round. After completion of the second round, a numeric code was assigned to each participant, allowing analysis of anonymized data. All data were handled and stored in compliance with the European General Data Protection Regulation.

### Identification of potential welfare indicators

Potential welfare indicators were sourced from a systematic literature review that aimed to collect valid and feasible welfare indicators for captive parrots^16^. The outcome measures identified in this systematic review were classified as animal-based indicators (e.g. excessive vocalization, stereotypies, responses to novel object or familiar and unfamiliar humans; n= 64), or environment-based indicators based on risk factors associated with an outcome measure (e.g. provision of foraging enrichment, social housing, cage size, diet composition, manual restraint; n= 35). Twenty-three additional animal- and environment-based welfare indicators were added after consultation with an avian medicine specialist (YvZ), resulting in a total of 122 potential welfare indicators presented to the participants, of which 79 were animal-based and 43 environment-based (Table S1, Table S2).

### Recruitment of participants

Participants were recruited by distributing a flyer containing a direct link to an online recruitment form and created with the software “LimeSurvey”. The flyer was distributed physically, shared through social media and online forums, and sent by email to potential participants. In the recruitment form, participants were asked to fill out their name and surname, type of expertise, years of experience working with parrots, contact email and professional or personal website. In order to increase our sample size, we also employed the snowball sampling method, meaning that people that registered to our survey could invite new participants by sharing the flyer with their working network^31^. There are no standards to select participants for the Delphi method, and various ways to qualify someone as “expert”^32^. Participants were selected on the basis of their expertise (veterinarian, researcher, behavioural consultant, animal keeper, breeder, other) and years of experience working with parrots (minimum 1 year experience required) as these are commonly accepted requirements in Delphi studies^32–34^. Being a parrot owner was not considered a sufficient type of expertise. One-hundred and fourteen participants that filled out the recruitment form passed the selection criteria and were invited to participate in the survey.

### Data collection

#### First round of the survey

This Delphi consultation consisted of two rounds of survey. The first round of survey was created using the online software “LimeSurvey” and it was divided in five sections.

The first section inquired participants about their demographic information: area of expertise (veterinarian, researcher, behavioural consultant, animal keeper, breeder, other), number of years of experience working with parrots (generally) and specifically with companion parrots, and current country of residence.

Before starting the second section, participants were invited to imagine the following scenario:

> “*You are invited to assess the welfare of a parrot kept as a companion animal. The parrot lives in a house with its owner(s). You are in the house, in front of the parrot and you have to consider the use of several measures in order to assess its welfare. The term ‘parrot’ refers to all species belonging to the order Psittaciformes*”.

In the second section, participants were presented with a list of 79 animal-based indicators grouped in separated categories according to commonalities in their underlying biological construct: abnormal and fear-related behaviours, exploratory behaviours, parrot-human interactions, locomotor behaviours, maintenance behaviours, social behaviours, sexual behaviours, body displays, and body measurements (table S1). At the end of each category, participants could add animal-based indicators that they considered important but that were missing from the list. For each indicator the participants needed to indicate whether they considered it a valid welfare indicator for all/most parrot species, valid only for certain species, or not valid as a welfare indicator. In the survey, we informed participants that “*we defined ‘welfare’ as the physical, physiological, and mental state of the parrot in relation to its environment and ‘valid welfare indicator’ a behavioural or physical measure that provides meaningful information about the welfare state of the parrot”*. As our aim was also to identify welfare indicators that could be used by owners, we also asked participants to indicate whether they considered taking the measure feasible for the parrot’s owner, feasible only for experts, or not feasible at all. In the present study we used the term ‘owner’ to refer to the person caring for the parrot, so we use this term as a synonym for ‘caretaker’, and not as the concept of owning a living being. In the survey, we defined *““feasible”* as *“a behavioural or physical measure that could be readily taken within 10 min, without causing acute stress responses of the parrot. This may include the use of minimally invasive routine handling techniques and/or commonly available equipment (e.g. weight scale)”*.

In the third section, the participants were presented with a list of 43 environment-based indicators grouped in five categories according to the husbandry or management condition that they represented: housing conditions, provision of enrichment, parrot-human interactions, nutrition, social needs (table S2). At the end of each category, participants could add environment-based indicators that they considered important but that were missing from the list. Participants were also asked to indicate to what extent the husbandry and management conditions had an impact on companion parrot welfare (high, medium, low) and whether these had an impact on welfare of all/most of the species or only on certain species. All environment-based indicators represented husbandry and management conditions that could be easily identified by owners (e.g. living alone vs in group, provision of enrichment, cage characteristics); therefore, we deemed it unnecessary to inquire participants about their feasibility.

In the fourth section, participants were asked to evaluate factors that could potentially affect parrot welfare. With the exception of one factor (hand-rearing with or without siblings), these factors were distilled from the scientific literature through a systematic review^16^, and subdivided into four categories: types of diet, early life/rearing history, species/sex/personality susceptibility to behaviour or medical problems, and species’ suitability to be kept as companion animal. These factors were presented as sentences to complete or statements and participants had to choose between three answer options: *always balanced, balanced but only for some species, always unbalanced* for the category types of diet; *less likely to develop/show welfare problems*, *more likely to develop/show welfare problems*, *neither more nor less likely to develop/show welfare problems* for the category early life/rearing history; *agree, neither agree nor disagree, disagree* for the categories species/sex/personality susceptibility to behaviour or medical problems, and species’ suitability to be kept as companion animal.

In the fifth section, participants were presented with a list of all animal- (n=79) and environment-based (n=43) indicators and asked to select and rank the 10 animal- and the 10 environment-based welfare indicators that they considered the most important for parrot welfare (1=most important; 10=least important).

Prior to sending out the survey, a pilot study was conducted to establish the clarity and appropriateness of the questionnaire, its structure, organization and items, and to determine the time required to complete it. Nine volunteers with an academic background in animal behaviour and welfare reviewed the pilot survey, reporting a completion time of approximately 40 minutes. Refinements were made according to the feedback of the participants of the pilot study.

The final survey was sent out for the first round and made available for five weeks (from May 18^th^, 2023 until June 22^nd^, 2023). Of the 114 participants that were invited to participate, 32 (28%) completed the entire survey and another 10 (8.8%) completed at least one category within the survey, which was our minimum requirement to be invited to the second survey round.

#### Second survey round

The second round of survey was created using Microsoft Word (365) and personalized for each participant to only include the items that were answered by this participant in the first survey round (see survey template in supplemental material).

The second survey contained the same sections and items presented in the first round of survey, except for the first one (demographic information). Participants were asked to review their answers based on the general agreement between all participants calculated for each parameter from the first round, which was presented to them in the respective sections. Participants could decide based on these results to either alter their previous answers or not. The participants were also presented with new animal- and environment-based indicators that had been suggested by one or more participants during the first survey round (Table S3, Table S4) and, similar to the first round, asked to assess these for their validity and feasibility (animal-based indicators) or for their impact on parrot welfare and applicability to all or most of the species (environment-based indicators).

In the third section, participants were asked to review their answers based on the general agreement between all participants calculated from the first round for the four categories of factors that could potentially affect welfare (i.e. diet, early life and pre-acquirement experiences, species and sex susceptibility to behaviour or medical problems, and species suitability as companion animal). Similar to the previous sections, participants could opt to alter their previous answers or leave these unchanged. When selecting the answer option “agree” on statements related to the species and sex susceptibility to behavioural or medical problems or suitability as companion, the participants were asked to review the lists of species or sex proposed by some participants in round one and to reply whether they agreed, disagreed, or neither agreed nor disagreed (see survey template in supplemental material).

In the last section, participants were presented with the list of the 10 animal-based and environment-based indicators that were deemed most important for parrot welfare according to the majority of the participants. The selection was made based on the rank score of each parameter, which was calculated as followed:

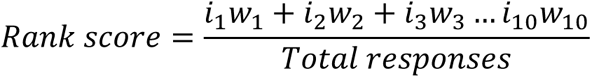

where:

*i*_*n*_= number of participants that selected the indicator in rank position n (1, 2, 3…10)

*w*_*n*_= weight on the rank position (e.g. n-rank position 1=10, n-rank position 10=1)

Participants were provided with the option to leave the ranking as it was, or re-rank the indicators if they disagreed with the presented order.

Prior to sending out the email with the second round of survey to all 42 participants that completed the first round, a pilot survey was again created to evaluate the second survey round in a similar manner as done for round one. This pilot was subsequently reviewed by seven of the nine academics who reviewed the first pilot. Following sending out of the second survey round (July 20^th^, 2023), participants were given nine weeks to complete the survey (by September 14^th^, 2023).

### Data analysis

All data collected from both the first and the second rounds of survey were analysed using descriptive statistics with the R statistical software^35^. For all items and for both rounds of survey, we calculated the percentage of participants that chose a specific answer option for a given item. While no standards exist to calculate consensus in Delphi studies^22, 36, 37^, consensus was estimated by calculating the percentage of agreement between participants (using the “dplyr” function in R^38^), following guidelines provided in previous Delphi studies on animal welfare indicators^23–25, 27^. Consensus was considered to have been achieved if an agreement of at least 70% was reached. Only those animal-based indicators for which consensus was reached for both the answer options “valid for all/most of the species” and “feasible for owners” were considered to be valid and feasible as indicators to assess the welfare of a companion parrot by caretakers. Similarly, environment-based indicators were considered valid and feasible to be used by caretakers if consensus was reached for both the answer options “high impact on welfare” and “applicable to/all most of the species”.

For the items in the third section on which consensus was reached for the answer option “agree”, data were further analysed by calculating the percentage of agreement between participants based on the species or the sex suggested to be more likely to develop specific behavioural problems, disease or pathology or the species suggested to be more or less suitable as companion animal.

Group stability between rounds was calculated by using the intraclass correlation coefficient (ICC) (R package “psych”^39^) as this is considered a reliable method to establish stability in Delphi studies^34^. All answer options were converted in scores from 1 to 3 and an ICC was computed for each section (animal-based indicators, environment-based indicators, factors that may impair welfare) by combining all answers submitted by the participants. The ICC value ranges between 0 and 1, with values between 0.75 and 0.90, and value greater than 0.90 representing good and excellent stability, respectively^40^. For the variable “applicability” (section environment-based indicators) it was not possible to calculate an ICC as this item was binomial (applicable to all/most of the species, applicable to only some species). For this parameter, stability was therefore established by calculating the percentage of times the group altered its answers from the first round to the second round and considered to be stable if changes occurred in less than 15% of cases, as previously suggested by Scheibe et al.^41^

To evaluate the potential effect of experience on the stability of the participants’ answers, we ran a generalized linear model that included proportion of times single participants altered their answers from the first to the second round as response variable and years of experience working with parrots as predictor.

Finally, to evaluate whether the rank of the most important animal- and environment-based indicators changed from the first to the second round, we calculated the rank score for each indicator by applying the formula as listed above to analyse the responses submitted in the first round. The rank scores from the first and the second round were then visually compared to verify if the indicators changed their position in the ranks.

## Results

### Demographics

A total of 42 participants from 14 countries (Austria, Brazil, Denmark, France, Germany, Indonesia, Italy, Pakistan, Spain, Sweden, Switzerland, the Netherlands, United Kingdom, United States of America) completed the first round of the survey. Twenty-one out of 42 participants (50%) completed the second round of survey. All five types of expertise were indicated by participants and experience working with parrots ranged from 1 to 51 years (Table 1). The percentage of participants with experience working with companion parrots was 85.7% in both rounds.

**Table 1.** Demographic information from the first and second round of survey.

|  | Area of expertise % (number of participants) |  |  |  |  |  |
| --- | --- | --- | --- | --- | --- | --- |
| Round | Veterinarian | Researcher | Breeder | Animal keeper | Behavioural consultant | Other |
| 1 | 47.6% (20) | 42.8% (18) | 9.5% (4) | 31% (13) | 19.0% (8) | bird curator (1), welfare organization president (1), sanctuary operator (1), zoologist (1) |
| 2 | 28.6% (6) | 57.7% (12) | 9.0% (2) | 28.6% (6) | 19.0% (4) | bird curator (1), welfare organization president (1) |
|  | Years of experience working with parrots % (number of participants) |  |  |  |  |  |
| Round | < 5 years | Between 5 and 10 years | Between 11 and 20 years |  | Between 21 and 51 years |  |
| 1 | 19% (8) | 14.4% (6) | 23.8% (10) |  | 42.8% (18) |  |
| 2 | 23.8 % (5) | 23.8 % (5) | 23.8 % (5) |  | 28.6% (6) |  |

### Group stability and influence of years of experience

All sections of the surveys showed excellent group stability between rounds: (animal-based indicators: ICC2k= 0.93; p<0.001; environment-based indicators: ICC2k= 0.94; p<0.001; factors impairing welfare: ICC2k= 0.94; p<0.001). The binomial variable “applicability” also achieved excellent group stability with only 2.8% responses changing between rounds. We found a statistically significant effect of years of working experience with parrots on the proportion of times that participants altered their responses between rounds, whereby the likelihood of an altered response decreased by 0.01 times with an increase in the number of years of experience (S.E.= 0.005722, z-value= −2.715, p= 0.007).

### Animal-based indicators

Thirty-two animal-based indicators (40.50%) reached consensus for being valid welfare indicators for all/most of the parrot species and feasible to be collected by owners (Table 2). Additionally, consensus was reached on five new animal-based indicators that were suggested during the first round (Table 2). These 37 animal-based indicators covered all nine welfare dimensions identified previously by Piseddu et al. (2024^16^; Table 2). Four animal-based indicators reached the consensus for being valid for all/most of the species but feasible only for experts whereas for the remaining answer options (“valid for only some species”, “not valid” and “not feasible”) no consensus was reached (Table S5).

**Table 2.**
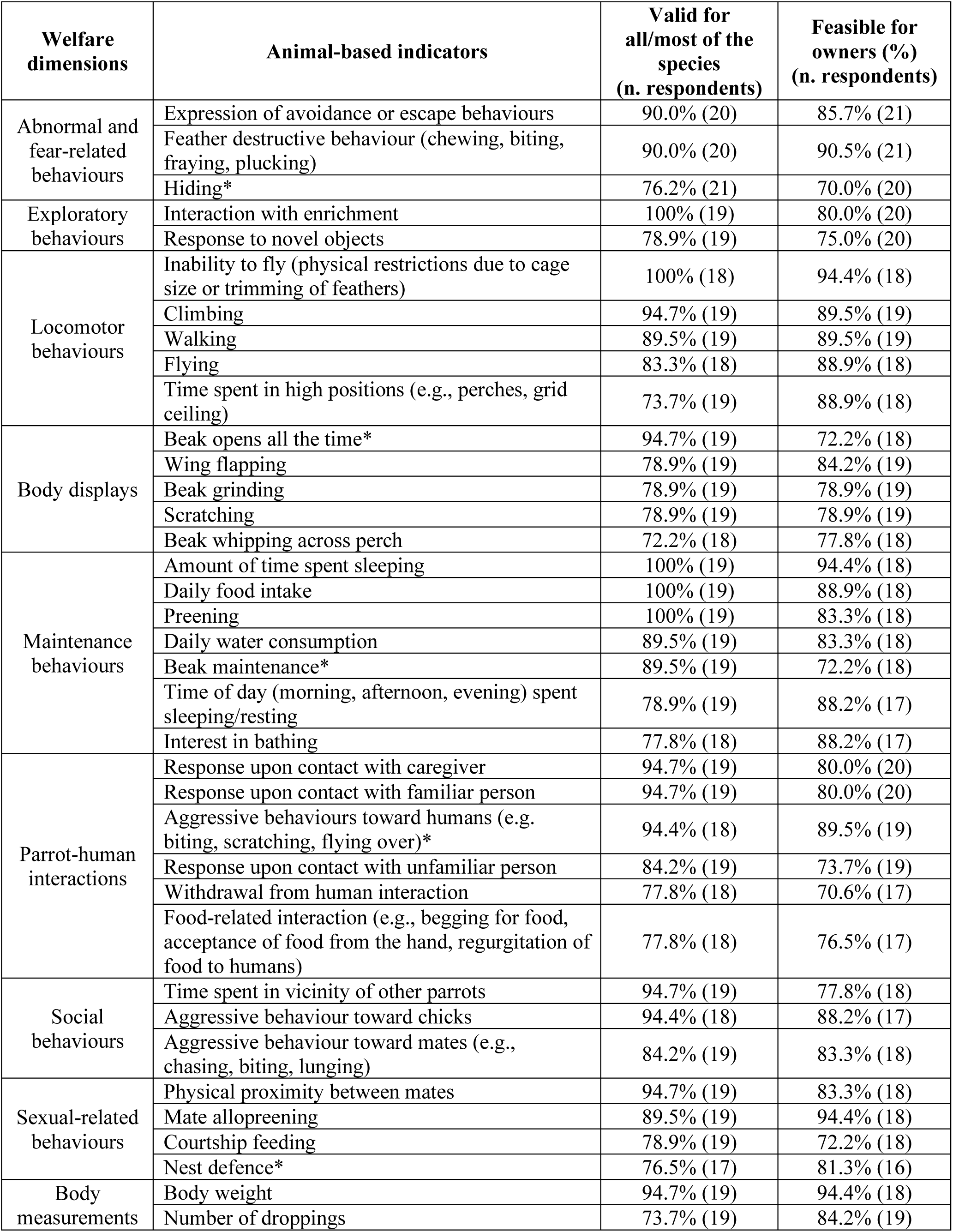
List of the animal-based indicators that reached the 70% agreement for both the answer options “valid for all/most of the species” and “feasible for owners”. Asterisk indicates animal-based indicators that were suggested by some participants in the first round of survey and therefore proposed only in the second round of the survey.

### Environment-based indicators

Twenty-six environment-based indicators (60.46%) reached consensus for having a high impact on parrot welfare and being applicable to all/most of the species (Table 3). Additionally, consensus was reached for ten new environment-based indicators that were proposed during the first round (Table 3). These 36 environment-based indicators covered all five categories of husbandry and management conditions as previously proposed in the systematic literature review^16^ (Table 3). Two indicators reached consensus for having moderate impact on welfare and being applicable to all/most of the species whereas for all remaining environment-based indicators, consensus was reached for the answer option “applicable to all/most of species” but no agreement was achieved about their impact on parrot welfare (Table 3, Table S6).

**Table 3.** List of the environment-based indicators that reached the 70% agreement for both the answer options “high impact on welfare” and “applicable to all/most of the species”. Asterisk indicates environment-based indicators that were suggested by some participants in the first round of survey and therefore proposed only in the second round of the survey.

| Husbandry and management conditions | Environment-based indicators | High impact on welfare (n. tot respondents) | Applicability to all/most of the species (n. tot respondents) |
| --- | --- | --- | --- |
| Housing | Cage characteristics (e.g., dimension, material, bars orientation) | 95.0% (20) | 100% (19) |
|  | Time spent out of the cage | 95.0% (20) | 90.0% (20) |
|  | Presence of a retreating area/room to rest, sleep or withdraw* | 90.0% (20) | 100% (20) |
|  | Provision of bathing opportunities | 90.0% (20) | 90.0 (20) |
|  | Air quality (e.g. presence of air purifier, exposure to fresh air)* | 85.0% (20) | 100% (20) |
|  | Exposure to direct sunlight/UV light* | 80.0% (20) | 100% (20) |
|  | Perches' characteristics (e.g., diameter, material) | 75.0% (20) | 100% (20) |
|  | Position and height of the perches (e.g., in relation to feeders or human-eye level) | 70.0% (20) | 100% (20) |
|  | Room where the cage is positioned (kitchen, living room, bedroom, etc.) | 70.0% (20) | 100% (20) |
|  | Frequency of cage cleaning* | 70.0% (20) | 100% (20) |
|  | Exposure to noise* | 70.0% (20) | 100% (20) |
|  | Environmental temperature | 70.0 % (20) | 90.0% (20) |
| Enrichment | Provision of cognitive enrichment | 100% (19) | 100% (19) |
|  | Variety of enrichment provided | 100% (20) | 100% (20) |
|  | Provision of chewable items | 100% (20) | 95.0% (20) |
|  | Provision of foraging enrichment | 95.0% (20) | 100% (20) |
|  | Opportunities to do physical exercise (flying, climbing, etc.) | 95.0% (20) | 95.0% (20) |
|  | Rotation of enrichment | 85.0% (20) | 100% (19) |
|  | Amount of enrichment provided | 75.0% (20) | 100% (19) |
|  | Opportunities to select items based on preference (e.g., for colour, shape or type of material) | 75.0% (20) | 100% (20) |
| Nutrition | Availability of clean fresh water* | 100% (20) | 100% (20) |
|  | Composition of the diet (quantity of fat, cholesterol, fibre, etc.) | 100% (20) | 95% (20) |
|  | Frequency of cleaning the food bowls* | 95.0% (20) | 100% (20) |
|  | Variety of food items provided | 85.0% (20) | 100% (20) |
|  | Manner/way in which food is offered to the bird (presented in a bowl, via enrichment, etc.) | 90.0% (20) | 100% (20) |
|  | Frequency with which food is provided | 78.9% (19%) | 100% (19) |
|  | Location and number of feeding areas* | 70.0% (20) | 100% (20) |
|  | Frequency of fresh food provision* | 70.0% (20) | 100% (20) |
| Parrot-Human interaction | Frequency/duration of manual restraint | 100% (20) | 100% (20) |
|  | Rearing history | 85.0% (20) | 100% (20) |
|  | Type of interaction with human (training, mouth to beak feeding, etc.) | 84.2% (19) | 100% (19) |
| Social needs | Level of social contacts (only vocal, visual and vocal, physical) | 100% (20) | 100% (18) |
|  | Frequency/duration of social separation events | 90.0% (20) | 100% (18) |
|  | Partner/cage mate choice (free vs imposed)* | 89.5% (19) | 94.7% (19) |
|  | Social housing (alone vs pair vs group) | 84.2% (19) | 100% (18) |
|  | Type of social companionship (same vs different in terms of species, size, sex, origin, etc.) | 75.0% (20) | 100% (18) |

### Rank

Abnormal and fear-related behaviours represented 5 of the top-6 ranking answers with feather destructive behaviour deemed as the most important indicator (Table 4). For all ten animal-based indicators included in the final ranking, consensus was reached for being valid indicators for all/most of the species, but four indicators did not reach consensus for being feasible for owners (Table 4). Among environment-based indicators, those related to provision of enrichment were the most recurrently chosen (n=4) with opportunities to do physical exercise and time spent out of the cage deemed as the most important ex-aequo. For all ten environment-based indicators included in the final ranking, consensus was reached for the answer option “applicable to all/most of the species”, whereas for the answer option “high impact on welfare”, consensus was reached for all except for the indicator “access to outdoor spaces” (Table 4). The ranking of both animal- and environment-based indicators did not change much between the two rounds, and mostly just one position, with exception of “level of activity” which moved up two ex aequo positions compared to round one (Table 4).

**Table 4.** Ranking of the 10 animal-based and environment-based indicators that were proposed by the participants as the most important indicators to assess parrot welfare. Asterisk indicates animal- and environment-based indicators for which consensus was not reached for the answer options “feasible for owners” and “high impact on welfare”, respectively.

| Welfare dimension | Animal-based indicators | Rank score round 2 | Rank position round 2 | Rank position round 1 |
| --- | --- | --- | --- | --- |
| Abnormal and fear-related behaviours | Feather destructive behaviour (chewing, biting, fraying, plucking) | 4.8 | 1 | 1 |
|  | Whole body stereotypies (head bobbing, rocking) * | 4.5 | 2 | 2 |
|  | Expression of avoidance or escape behaviours | 3.8 | 3 | 4 |
|  | Locomotor stereotypies (route-tracing, pacing) * | 3.7 | 4 | 4 |
| Locomotor behaviour | Level of activity (time spent inactive vs active) * | 3.5 | 5 | 3 |
| Abnormal and fear-related behaviour | Excessive vocalization/screaming* | 2.7 | 6 | 6 |
| Exploratory behaviour | Interaction with enrichment | 2.2 | 7 | 8 |
| Maintenance behaviour | Daily food intake | 2.1 | 8 | 8 |
| Locomotor behaviour | Inability to fly (physical restrictions due to cage size or trimming of feathers) | 1.7 | 9 | 9 |
| Parrot-human interaction | Response upon contact with caregiver | 1.0 | 10 | 10 |
| Husbandry and management conditions | Environment-based indicators | Rank score round 2 | Rank position round 2 | Rank position round 1 |
| Enrichment | Opportunities to do physical exercise (flying, climbing, etc.) | 4.5 | 1 | 2 |
| Housing | Time spent out of the cage | 4.5 | 1 | 1 |
|  | Cage characteristics (e.g., dimension, material, bars orientation) | 4.0 | 3 | 3 |
| Enrichment | Provision of foraging enrichment | 3.5 | 4 | 5 |
| Social need | Social housing (alone vs pair vs group) | 3.4 | 5 | 4 |
| Enrichment | Provision of cognitive enrichment | 3.0 | 6 | 6 |
| Nutrition | Composition of the diet (quantity of fat, cholesterol, fibre, etc.) | 2.7 | 7 | 7 |
| Parrot-human interaction | Rearing history | 1.9 | 8 | 8 |
| Enrichment | Variety of enrichment provided | 1.7 | 9 | 9 |
| Housing | Access to outdoor spaces* | 1.0 | 10 | 10 |

### Factors with an impact on parrot welfare

#### Type of diet

Expert consultation revealed consensus for diets always being unbalanced if exclusively based on seeds (89.5%), based on only one type of seed (100%), or exclusively based on pellets (70.6%) (Table S7). Regarding diets exclusively based on mashed food, no consensus was reached (Table S7).

#### Early-life and pre-acquirement experiences

Participants reached consensus on the statements that hand-reared parrots, wild-caught parrots and parrots acquired before the end of weaning are more likely to develop/show welfare problems. Additionally, consensus was reached for the statement that parrots that are hand-reared with siblings (vs hand-reared alone) are less likely to develop/show welfare problems (Table S8). For all other remaining statements, no consensus was reached (Table S8).

#### Sex and species susceptibility to develop behavioural problems, diseases or pathological conditions

Participants reached consensus for the statement that “*some parrot species are more likely to develop behavioural problems when kept in captivity*” (83% agreement; Table S9). Participants agreed that cockatoos (excluding cockatiels) and grey parrots (*Psittacus erithacus*) are more likely to develop behavioural problems when kept in captivity (93.3% and 86.7% agreement, respectively; Table S10). Consensus was also reached on cockatoos (excluding cockatiels) being at greater risk for developing feather damaging behaviour, aggressiveness, and hormonal behaviours (100%, 92.8% and 100% agreement, respectively; Table S11), while for grey parrots agreement was reached only for feather damaging behaviour (100% agreement, Table S11). For the remaining 22 species that were suggested to be prone to develop behaviour problems in captivity, no group consensus was reached for any of the answer options provided (Table S10).

#### Personality and suitability of species as companion parrots

A 100% consensus was reached on the statement that “*assessing parrot personality can improve/ensure parrot welfare*”, while 75% of the participants agreed with the statements “*some parrot species are more suitable to be kept as companion animals*” and “*some parrot species should not be kept as companion animals*” (Table S9). Participants agreed for 7 out of 28 species/genera that these would be more suitable to be kept as companion animals: lovebirds (*Agapornis* spp.; 86.7% agreement, n= 15), budgerigars (*Melopsittacus undulatus*; 100.0% agreement, n= 15), *Pyrrhura* spp. (76.9% agreement, n= 13), green-cheeked conure (*Pyrrhura molinae*) (76.9% agreement, n= 13), cockatiel (*Nymphicus hollandicus;* 93.3% agreement, n= 15), parrotlets (*Forpus* spp.; 78.6% agreement, n= 14), and monk parakeets (*Myiopsitta monachus*; 76.9% agreement, n= 13; Table S12). Conversely, participants (n=14) disagreed with the statement that white cockatoos (*Cacatua alba,* 92.9%) and long-billed corellas (*Cacatua tenuirostris,* 78.6%) would be suitable to be kept as companion animals. Finally, 76.9% of participants indicated to neither agree nor disagree with medium-sized species being suitable as companion animals (Table S12). Among participants, consensus was also reached that cockatoos (excluding cockatiels), large cockatoos, critically endangered species, and heavily trafficked species should not be kept as companion animals (agreement of 86.7%, n=15; 85.7%, n= 14; 75%, n= 16; and 81.3%, n= 16, respectively; Table S13).

## Discussion

### Demographic and stability of responses between rounds

Despite informing participants about the iterative nature of the project and making them feel engaged and actively involved to minimize the risk of dropping out^32, 34^, we obtained an attrition rate of almost 50% between rounds from our initial 41 participants. This might be due to the high number of items included in the survey^42^, which may also explain why fewer participants filled out the survey compared to other Delphi studies on animal welfare that included fewer items ^23–25, 29^. Moreover, the field of parrot welfare is comparatively smaller than the field of the other species addressed in Delphi studies, potentially explaining the smaller panel size. There are currently no standards on the number of panellists required to obtain robust results, although our final panel size (n=21) was still in the range considered ideal (between 8 and 23 participants)^32^ for Delphi studies. Although small, our panel was comprised of participant with various types of expertise (academics, behavioural consultants, veterinarians, breeders, animal keepers) thereby being heterogeneous, which is generally recommended as it allows for multi-angled analysis of complex topics such as animal welfare, thereby leading to an increased quality of the results^43, 44^. Moreover, the number of participants was balanced between types of expertise, and this remained stable between rounds, thereby contributing to a high reliability of the results.

We found a relationship between years of work experience with parrots and the proportion of times that participants altered their responses between rounds, suggesting that the opinion of less experienced participants may be more influenced by seeing the results from the first round than participants with more experience. However, this effect was not strong enough to affect group stability, as participants were resolute on their answers and did not significantly change these at group level between rounds. The optimal number of rounds of survey in Delphi studies is considered to be three^34^; however, given that we had already achieved stability after the second round and considering the high attrition rate, we decided not to conduct an additional round.

### Indicators

We identified 32 animal-based indicators that reached consensus between participants for being considered valid for all/most of the parrot species and feasible for owners. These were different types of behaviours (i.e. abnormal, fear-related, locomotor, exploratory, social, human-directed, maintenance, sexual), body displays and body measurements. Abnormal and fear-related behaviours represented half of the ten most important animal-based indicators and occupied the highest positions in the rank. Three of these indicators did not reach consensus for being feasible for owners: whole body and locomotor stereotypies, and excessive screaming/vocalization. Stereotypies can reflect difficulties in the ability of an animal to cope with its environment^45^, which may explain why stereotypies were rated by participants among the most important animal-based indicators. However, stereotypies can be misinterpreted and difficult to recognize as abnormal behaviours for unexperienced owners, and are sometimes only performed when the animal is alone^5^. Moreover, abnormal repetitive behaviours can persist as ‘behavioural scars’ despite the triggering event or situation long being resolved^46^. As a result, a level of expertise is required for assessors to ascertain whether stereotypies are reflecting the parrot’s present welfare state. Similarly, excessive vocalizations and screaming are considered signs of poor welfare^5^, more specifically as behavioural responses to frustration, fear or lack of attention^47^. However, these again may be challenging for owners to distinguish from normal vocalizations as parrots are highly vocal species that also produce frequent and loud vocalization in a social context^8^. Level of activity was also included in the top-10 ranking of welfare indicators but considered by participants as not feasible for owners, most likely because evaluating parrot activity level would be highly time consuming. Many other indicators not included in the final rank were also considered valid, but also failed to reach consensus for feasibility. Mostly, this involved strong discordance between participants on whether the indicator was “feasible for owners” or “feasible, but only for experts”, which may result from differences in expert opinion or difficulties with accurate interpretation of this parameter through the chosen method of consultation. Possibly, experimental studies could provide a more effective strategy to assess the actual feasibility for owners to evaluate these welfare indicators.

The remaining animal-based indicators in the top 10 rank (feather destructive behaviours, expression of avoidance or escape behaviours, interaction with enrichment, daily food intake, inability to fly and response upon contact with caregiver) all reached consensus for being considered feasible for owners and, due to their importance, represent solid measurement for assessing parrot welfare. These body and behavioural measurements would therefore be a good starting point for owners to evaluate their bird’s welfare, which – together with the support of experts or new technologies^48^ – could produce a better overview of the parrot’s current welfare state.

All 43 environment-based indicators presented to participants reached a consensus for the answer “applicable to all/most of the species”, but only 26 indicators reached consensus for being considered as having a high impact of welfare. Opportunities for physical activities and time spent out of the cage were ranked by participants as the most important environment-based indicators, sharing the joint first place in the ranking, followed by cage characteristics and provision of foraging enrichment. Environmental enrichment and housing thus emerged as aspects that participants considered particularly important, likely because these provide parrots with an opportunity to meet their biological needs: mental stimulation, opportunities to forage and exercise, and interaction with other parrots - preferably conspecifics^4, 5, 7, 10, 49^. These 26 indicators would be valuable to inform owners about the type of husbandry and management conditions to provide and, if necessary, how to modify the current living environment. In addition, these indicators can also be used by professionals that works with companion parrots and shared in brochures with informative and educational purposes. For 16 indicators, opinions of the participants were highly split between the options “high” and “moderate impact” on parrot welfare. These results suggest that participants were more divided with regard to the severity of the impact, which might be related to the subjective nature of this parameter.

In the first survey round, participants suggested 46 new indicators (including 27 animal-based and 19 environment-based ones) as useful for evaluating parrot welfare in addition to those that had previously been identified through the systematic literature review. After the second round, consensus was reached on 14 of these (including 9 environment-based and 5 animal-based indicators). Although these indicators were not processed following the Delphi methodology due to being included only in the second round, the consensus reached in the second round still suggests that these indicators could be valuable and need to be taken into consideration as potential welfare indicators.

Several participants mentioned that the value of many indicators proposed highly depends on three aspects that were not taken into consideration in the survey: 1) the context in which a behaviour is observed or a measurement is taken (e.g. aggressive behaviour towards a human could indicate poor welfare in case of forced physical interactions or excessively fearful birds, but could also manifest itself as a natural behaviour to defend the partner or nest site during the breeding season)^50^; 2) the valence of the parameter, which could be positive or negative dependent on intensity, frequency, or duration (e.g. preening, time spent sleeping) or even on species (e.g. response to novel objects for neophilic vs. neophobic parrots^51^) and requires further validation due to being primarily based on anecdotal, non-scientific information (e.g. body displays such as beak grinding and wing flapping that are suggested to reflect contentment^52^ and threat display^10^, respectively); and 3) the personality of the individual, as all participants agreed that assessing parrots’ personality can benefit parrot welfare and studies have shown that specific individual traits (e.g. being neophobic, neurotic, explorative, proactive, vigilant) can influence parrots’ response to the provision of enrichment or food ^53, 54^, interactions with humans^55^, pairing success^56^, or the risk of developing abnormal behaviours^57, 58^. Therefore, it would be important to further research the effects of these factors on parrots.

### Factors with an impact on parrot welfare

We presented participants with factors that may impact companion parrot welfare, according to common knowledge or previous experimental studies. Participants concurred with most of these statements that are supported by scientific evidence, reaching consensus for 11 out of 17 statements, thereby further validating the impact of diet^6, 59, 60^, hand-rearing^59, 61–63^, of being wild-caught^59, 64^ and acquisition before weaning^59, 61^ on parrot welfare. Several of these factors, particularly those related to acquisition or rearing, are difficult or even impossible to alter, but could nevertheless still guide a prospective parrot’s owner in choosing the right companion parrot, thus reducing the risks of experiencing poor welfare.

Participants agreed that some species are more likely to develop behavioural problems when kept in captivity. Specifically cockatoos and grey parrots were evaluated by the majority of participants as species that are predisposed to develop feather damaging behaviour, which is in line with the literature^65–, 67^ and findings from other surveys^66–68^. Participants furthermore agreed that cockatoos are more likely to display aggressive and hormonal behaviours. Hormonal behaviours, sometimes referred to as sexual-related behaviours, are innate natural behaviours that parrots display during the breeding season (e.g. defending the nest site and their mate)^69^. In captivity, specific environmental cues (e.g. providing a nest and/or nesting material, high-fat diets) and physical interactions that unintentionally simulate courtships behaviours (e.g. touching the tail and back) are hypothesized to encourage parrots, especially hand-reared individuals, to pair-bond with one family member and display sexual-related behaviour toward that person^47, 69, 70^. These behaviours may include regurgitating food or attempting to copulate with the bonded human, masturbation, or aggression towards any potential rival that approaches its “mate”^47, 50, 71^, which may sometimes lead to redirected aggression towards the bonded human^50^. In accordance with this, Tygesen and Forkman found that emotional closeness as reported by owners and the frequency of interactions with their parrot were positively correlated with their parrot being aggressive towards humans^1^, in line with the problem of “hormonal behaviour”^47, 69^. These observations indicate that hormonal behaviours are of particular interest when keeping parrots in captivity, and likely require scientific investigation to determine their causes and consequences for parrot welfare and on the parrot-human relationship.

We found inconsistencies between participants’ opinion and the scientific literature for other aspects such as the benefit of neonatal handling^72–74^, species or sex susceptibility for certain diseases^6, 75–77^ or behavioural problems^61, 78–82^, and acquisition of parrots from pet shops or shelters^60, 66, 79^. These inconsistencies could be explained by the heterogeneity of expertise in our panel or other unidentified reasons. For example, topics such as species’ or gender predispositions to diseases or effects of neonatal handling may have required specific knowledge or higher levels of specialization^34^, thereby hindering some of the participants in their assessment.

### Suitability of parrots as companion animals

The statement “parrots should not be kept as companion animals” did not reach a consensus for any answer option available, and in fact revealed divided opinions across the three available answer options. This could be due to different ethical positions between participants on keeping parrots as companions. Participants agreed that “some parrots species should not be kept as companion animal” whereby cockatoos (excluding cockatiels) represented the only taxonomic group for which consensus was reached that these are not being suitable companion parrots. This is likely related to the already discussed sensitivity of these species to develop behavioural problems. Cockatoos are well known to be “difficult pets”; however, experimental studies on the welfare of these species in captivity are scarce, and mainly focused on feather damaging behaviour^16^. Consensus was also reached on heavily trafficked and critically endangered species^83^ being unsuitable companion animals. Parrots are among the most threatened species^84^ and poaching and illegal pet trade are considered two of the main causes^85^. Therefore, participants may have considered banning these species from the pet trade as a means to protect wild parrots’ populations from further decline.

Seven small-sized species reached consensus for being more suitable to be kept as companion animals. Participants may have perceived the species-typical needs for these species to be more easily fulfilled (e.g. social housing) than for others, hence making these species easier to be kept as companion animals. Many of these species possess biological characteristics (i.e. relative small brain size, foraging style that does not required extensive food handling) that were found to be linked to a lower likelihood to develop feather damaging behaviour and stereotypies when kept as companion animals^65^. Nevertheless, even if these seven species are perceived as more suitable companions, all of them, budgerigars^80, 81, 86, 87^, cockatiels^88–91^, conures^67^, lovebirds^63, 67^, pacific parrotlets (*Forpus coelestis*)^67^ and monk parakeets^6^, can show signs of poor welfare when kept in inappropriate conditions. As the results regarding specific species’ suitability as a companion animal were not processed in multiple rounds, further research should be performed to validate these findings, and evaluate in further detail whether and which biological characteristics would render a species more or less adaptable to the domestic environment.

### Conclusion

Using the Delphi method, we identified 37 animal-based and 36 environment-based welfare indicators that were evaluated by participants with expertise in parrot welfare as valid and feasible for parrot owners to assess the welfare of all/most parrot species. The expert panel also concurred with scientific findings regarding factors potentially affecting companion parrot welfare: types of diet, pre-acquirement experiences, species susceptibility to develop behavioural problems, and suitability of different species as companion animals. This science-based information could be used by parrot owners, (veterinary) professionals, and policy makers to monitor parrot welfare and improve husbandry conditions of captive parrots.

## Acknowledgments

We express our sincere gratitude to all participants of both first and second round of the survey that made the completion of this study possible by sharing their expertise and knowledge. Additionally, we would like to thank the colleagues of the Centre for Animal Nutrition and Welfare for having provided their feedback on the conceptualization and methodology of this study, all researchers that participated in both pilots and Stephanie Lürzel and Remco Folkertsma for their assistance in the statistical analysis.

## Author Contributions

Conceptualization, all authors; methodology, all authors; data collection, A.P.; statistical analysis, A.P; data curation, A.P.; writing—original draft preparation, A.P.; writing—review and editing A.P, Y.R.A.v.Z., J.-L. R.; supervision, Y.R.A.v.Z., J.-L. R.

## Data availability

All data generated or analysed during this study are included in this published article (and its Supplementary Information files)

