## Supplemental Information for "Evaluation of welfare indicators for companion parrots: a Delphi consultation survey"

### Methods

**Table S1**. List of all animal-based indicators presented to participants, grouped in welfare dimensions according to commonalities in their underlying biological construct. The asterisk indicates the animal-based indicators identified through consultation with an avian medicine specialist. The superscript [a] indicates animal-based indicators that were created combing more outcome measures obtained through the systematic review or/and after the consultation of the avian medicine specialist.

| Welfare dimension | Indicators |
|---|---|
| Abnormal and fear-related behaviours | Expression of avoidance or escape behaviours<br>Excessive vocalization/screaming<br>Feather destructive behaviour (chewing, biting, fraying, plucking)<br>Toe/nail biting*<br>Excessive chewing (e.g., wires)<br>Sham behaviours (e.g., sham chewing, sham flying, sham bathing)[a]<br>Whole body stereotypies (head bobbing, rocking)<br>Locomotor stereotypies (route-tracing, pacing) |
| Explorative behaviours | Response to novel objects<br>Response to novel food items<br>Interaction with enrichment<br>Time spent foraging<br>Response in unfamiliar environments |
| Locomotor behaviours | Level of activity (time spent inactive vs active)<br>Flying<br>Walking<br>Climbing<br>Hopping<br>Time spent on the ground/bottom of the cage<br>Time spent in high positions (e.g., perches, grid ceiling)<br>Inability to fly (physical restrictions due to cage size or trimming of feathers) |
| Maintenance behaviours | Amount of time spent sleeping<br>Time of day (morning, afternoon, evening) spent sleeping/resting<br>Posture while sleeping (e.g., on one leg, head tucked under wings)*<br>Daily food intake<br>Daily water consumption<br>Interest in bathing<br>Preening activity (incl. time of day and frequency, duration)[a] |
| Parrot-Human interactions | Response upon contact with caregiver<br>Response upon contact with familiar person<br>Response upon contact with unfamiliar person<br>Facial expressions (e.g., blushing) during human interaction<br>Ruffling of feathers (e.g., nape, crown, beard) during human interaction<br>Willingness to approach or step up on the human<br>Withdrawal from human interaction |

| | |
|---|---|
| | Food-related interaction (e.g., begging for food, acceptance of food from the hand, regurgitation of food to humans)[a]<br>Sexual related behaviours (e.g., panting, receptive posture)[a]<br>Behaviour during manual restraint (vocalizing, biting, resistance)<br>Contact seeking behaviours in absence of humans (e.g., vocalization, flapping wings)<br>Preference for some humans over others<br>Initiation of contact with human being<br>Mimicry of human sounds |
| Social behaviours | Aggressive behaviour toward non-mates (e.g., chasing, biting, lunging)<br>Aggressive behaviour toward mates (e.g., chasing, biting, lunging)<br>Aggressive behaviour toward chicks<br>Allopreening<br>Time spent in vicinity of other parrots<br>Frequency and duration of social interaction<br>Intraspecies aggression during feeding (incl. stealing of food)<br>Play behaviour towards cage-mates*<br>Vocal communication with other birds (e.g., type and frequency)[a] |
| Sexual behaviours | Courtship feeding<br>Physical proximity between mates<br>Mate allopreening<br>Copulation |
| Body displays | Nape and/or crown feather ruffling<br>Cheek or beard feather ruffling<br>Wing and/or leg stretch<br>Scratching<br>Beak grinding<br>Beak whipping across perch*<br>Wing flapping*<br>Toe/nail biting*<br>Tail wagging<br>Ruffling of body feathers<br>Erection of crest feathers |
| Body measurements | Body weight<br>Pectoral muscle condition score*<br>Condition of body feathers (contour/down feathers)<br>Condition of flight feathers (wing and tail)<br>Beak appearance (e.g., length, shape, position)*<br>Cere/nare appearance (e.g. colour, shape, size)*<br>Eye appearance (e.g., half-open, discharge)*<br>Mentation/alertness*<br>Posture (e.g., fluffed appearance, weight bearing)*<br>Gait changes (incl. flight)*<br>Respiration/breathing changes (e.g., frequency, depth)<br>Number of droppings*<br>Appearance of droppings (e.g., colour, consistency)* |

**Table S2.** List of all environment-based indicators presented to participants, grouped according to the husbandry or management condition. The asterisk indicates the environment-based indicators identified through consultation with an avian medicine specialist. The superscript [a] indicates environment-based indicators that were created combing more outcome measures obtained through the systematic review or/and after the consultation of the avian medicine specialist.

| **Husbandry and management condition** | **Indicators** |
|---|---|
| Housing | Cage characteristics (e.g., dimension, material, bars orientation)[a]<br>Environmental temperature<br>Perches' characteristics (e.g., diameter, material)[a]<br>Position and height of the perches (e.g., in relation to feeders or human-eye level)[a]<br>Time spent out of the cage<br>Position of the cage in the room<br>Room where the cage is positioned (kitchen, living room, bedroom, etc.)*<br>Exposure to artificial light at night<br>Access to outdoor spaces<br>Provision of bathing opportunities |
| Nutrition | Frequency with which food is provided<br>Time of day (morning, afternoon, evening) at which food is provided<br>Variety of food items provided |

| | |
|---|---|
| | Composition of the diet (quantity of fat, cholesterol, fibre, etc.)[a]<br>Manner/way in which food is offered to the bird (presented in a bowl, via enrichment, etc.)*<br>Provision of supplements (multivitamin, calcium, essential fatty acids, etc.)[a]<br>Frequency of cleaning the food bowls*<br>Pellet Size<br>Consumption of human food |
| Enrichment | Provision of cognitive enrichment<br>Provision of foraging enrichment<br>Provision of auditory enrichments*<br>Provision of visual enrichment*<br>Rotation of enrichment*<br>Variety of enrichment provided*<br>Amount of enrichment provided<br>Opportunities to do physical exercise (flying, climbing, etc.)[a]<br>Provision of chewable items<br>Provision of climbing toys, swings, and ladders<br>Opportunities to select items based on preference (e.g., for colour, shape or type of material)[a] |
| Parrot-human interactions | Time spent without presence of a human<br>Time spent on interaction with human<br>Type of interaction with human (training, mouth to beak feeding, etc.)[a]<br>Human produces loud noises and/or sudden movements<br>Frequency/duration of manual restraint<br>Number of people in the household<br>Number of people in the household that regularly interact with the parrot<br>Rearing history |
| Social needs | Social housing (alone vs pair vs group)<br>Opportunities for pair bonding (i.e., living with/without a mate)<br>Type of social companionship (same vs different in terms of species, size, sex, origin, etc.)[a]<br>Frequency/duration of social separation events<br>Level of social contacts (only vocal, visual and vocal, physical)* |

**Table S3**. List of all animal-based indicators presented to participants in the first round of survey, grouped in welfare dimensions according to commonalities in their underlying biological construct.

| **Welfare dimension** | **Indicators proposed by the experts in round 1** |
|---|---|
| Abnormal and fear-related behaviours | Hiding<br>Excessive masturbation<br>Tonic-clonic immobility/freezing<br>Grunting<br>Tremors<br>Spot pecking |
| Explorative behaviours | Response to electronic devices (TV, music...) |
| Locomotor behaviours | Swinging |
| Maintenance behaviours | Beak maintenance |
| Parrot-Human interactions | Aggression towards humans linked to a specific location or perimeter<br>Eating when they see the humans eating<br>Expression of abnormal behaviours (e.g. toe/nail bite, feather damaging behaviours) when human is present<br>Aggressive behaviours toward humans (e.g. biting, scratching, flying over) |
| Social behaviours | Displacement behaviours during social interactions |
| Sexual behaviours | Nest building<br>Nest defence<br>Masturbation<br>Destruction of nesting materials<br>Searching for nesting areas<br>Territoriality during breeding |
| Body displays | Beak open all the time<br>Absence of any type of body display<br>Pupil movements |
| Body measurements | Respiratory effort at rest<br>Respiratory effort when disturbed<br>Prolapses<br>Length and frequency of moulting |

**Table S4**. List of environment-based indicators presented to the participants in the first round of survey grouped according to the husbandry or management condition.

| Husbandry and management condition | Indicators proposed by the experts in round 1 |
|---|---|
| Housing | Size of the room where the cage is positioned<br>Environmental humidity<br>Presence of a retreating area/room to rest, sleep or withdraw<br>Presence of a platform to stand on<br>Size/number of windows in the room<br>Presence of a nesting area<br>Frequency of cage cleaning<br>Exposure to noise<br>Air quality (e.g. presence of air purifier, exposure to fresh air)<br>Exposure to direct sunlight/UV light<br>Artificial light characteristics (e.g. type, intensity) |
| Nutrition | Balance between provision of fresh and dried food<br>Location and number of feeding areas<br>Way in which food is stored<br>Availability of clean fresh water<br>Frequency of fresh food provision |
| Enrichment | Size of enrichment in relation to parrot size<br>Area of the room/cage where enrichment is placed |
| Social needs | Partner/cage mate choice (free vs imposed) |

### Template of the second round of survey

#### Information and Consent

Dear Expert,

Thank you for taking part in the second and last round of the survey. The aim of the second survey is for you to review the general agreement between experts on answers submitted in the previous survey, and decide whether or not you want to alter your previous responses. In addition, you can share your opinion regarding new information proposed by experts in the previous round.

Please remember to save your changes regularly while you are filling out the document to ensure that your answers are stored.

During the first survey, many of you noted that several indicators presented in the lists are relevant for parrot welfare, but only in specific circumstances and contexts. We have not yet included these refinements in the survey as we acknowledge that everyone's time is precious and limited, and would like to ensure that as many people as possible will complete the survey. Therefore, additional aspects such as age, species, previous experiences, personality, and also the valence of the indicators (positive/negative) will be evaluated at a later time, through an expert panel consultation. If you would like to take part in this expert panel consultation, please tick the following box:

**Yes, I would like to be contacted in the future to participate in the expert panel consultation** ☐

**Please note that:**

• Participation is voluntary and anonymous.

• You can withdraw at any moment of the study.

• You have to be at least 18 years old to participate.

• The results of this survey will be included in a PhD thesis and published in a scientific journal.

• Your name will not be visible for the other participants, it will not appear in any published documents related to this study, and your statements will remain anonymous.

• Only the responsible person of this study will be able to trace back your identity and corresponding responses.

• Your personal data will be treated in compliance with the European General Data Protection Regulation and will be stored only as long as needed for the purpose of this study.

Before starting to fill out the questionnaire please tick the box below:
**I confirm to have read and understood the above statements and I give my informed consent.** ☐

As a measure of appreciation, if you could send us a postal address after completing this survey, we will ship you a small gift to thank you.

If you have any questions or concerns, please do not hesitate to contact the responsible person of this study:

**Andrea Piseddu**
Institute of Animal Welfare Science, University of Veterinary Medicine Vienna,
Vienna, Veterinaerplatz 1, 1210 Vienna, Austria


#### Indicators

In the next two sections you will be presented with the list of animal-based measures (behaviours and physical measures) that you already assessed for their validity and feasibility, and the list of environment-based measures (husbandry and management conditions) that you already evaluated for their influence on companion parrot welfare as well as for their applicability to all/most of the species or only to some parrot species. This time you will also see in bold the percentage of experts that selected the available options (e.g. **80%**) and your previous responses indicated as ticked boxes (☒) from the first round of the survey. In this second round you have the opportunity to review the general agreement between experts and decide whether or not you want to alter your previous responses. If you decide to alter your response, please deselect the ticked box and tick the box next to your new answer.

In this second round, you can also assess the validity, feasibility, impact on welfare and applicability of new animal- and environment-based indicators proposed by you or other experts in the first round.

**Please keep in mind the following information while you are answering:**

-Scenario: you are invited to assess the welfare of *a parrot kept as a companion animal*. The parrot lives in a house with its owner(s). You are in the house, in front of the parrot and you have to consider the use of several measures in order to assess its welfare.

-Welfare: physical, physiological, and mental state of the parrot in relation to its environment.

-Valid welfare indicator: behavioural or physical measure that provides meaningful information about the welfare state of the parrot.

-Feasible indicator: behavioural or physical measure that can be readily taken within 10 min, without causing acute stress responses of the parrot. This may include the use of minimally invasive routine handling techniques and/or commonly available equipment (for example weight scale).

-The term "parrot" refers to all species belonging to the order *Psittaciformes*.

-We define "cockatoos" as all the species belonging to the superfamily *Cacatuoidea*, excluding cockatiels (*Nymphicus hollandicus*). White cockatoo species (e.g. *C. sulphurea*, *C. alba*, *C. galerita*, *C. moluccensis*,) can be indicated with "*Cacatua* spp.".

We define "macaws" as the species belonging to the genera *Ara*, *Anodorhynchus*, *Cyanopsitta*, *Orthopsittaca*, *Primolius* and *Diopsittaca*.

##### Animal-based indicators

For the option "Valid, but only for some species" you can find the list of species/taxonomic groups for which some experts proposed the measure as a valid indicator of welfare, followed by the number of experts that indicated them (e.g. *Amazona* spp.= indicated by one expert, *Amazona* spp. (2) = indicated by two experts). You can modify the list by adding or deleting species/taxonomic groups. Please modify the list **only** if you select the option "Valid, but only for some species".

**Please select** the option "Valid, but only for some species" if the measure could be used as indicator of welfare:

- only in some species or taxonomic groups, regardless of the context

**Please do not select** the option "Valid, but only for some species" if the measure could be used as indicator of welfare in all/most of the species:

- but only in specific contexts.
- even if particularly important for only some species
- except for a few of them.

Instead, please select the option "Valid for all/most of the species"

| | Validity | | | | Feasibility | | |
|---|---|---|---|---|---|---|---|
| **Abnormal and fear related behaviours** | Valid for all/most of the species: | Valid, but only for some species | | Not valid | Feasible for owners | Feasible, but only for experts | Not feasible |
| Expression of avoidance or escape behaviours | **80%** ☐ | **7.5%** ☐ | **Valid only for:**<br>*Amazona* spp.<br>*Anodorhynchus* spp.<br>*Ara* spp.<br>*Psittacus erithacus* | **12.5%** ☐ | **77%** ☐ | **20%** ☐ | **3%** ☐ |
| Excessive vocalization/screaming | **62.5%** ☐ | **30%** ☐ | **Valid only for:**<br>*Amazona* spp. (5)<br>Macaws (3)<br>*Ara* spp.<br>Cockatoos (2)<br>*Cacatua alba*<br>*Cacatua galerita*<br>*Cacatua ophtalmica*<br>Larger parrots (*2*)<br>Parakeets<br>*Poichephalus* spp.<br>*Psittacus Erithacus* (*4*) | **7.5%** ☐ | **74%** ☐ | **23%** ☐ | **3%** ☐ |
| Feather destructive behaviour (chewing, biting, fraying, plucking) | **85%** ☐ | **10%** ☐ | **Valid only for:**<br>*Cacatua alba*<br>*Cacatua galerita*<br>*Cacatua ophtalmica*<br>Larger parrots<br>*Psittacus erithacus*<br>*Trichoglossus haematodus*<br>*Trichoglossus moluccanus* | **5%** ☐ | **90%** ☐ | **5%** ☐ | **5%** ☐ |
| Toe/nail biting | **73%** ☐ | **19%** ☐ | **Valid only for:**<br>*Anodorhynchus* spp.<br>*Ara* spp.<br>*Nymphicus hollandicus*<br>*Eclectus roratus*<br>*Eolophus roseicapilla*<br>*Psittacus Erithacus* (2) | **8%** ☐ | **71%** ☐ | **26%** ☐ | **3%** ☐ |
| Excessive chewing (e.g., wires) | **59%** ☐ | **18%** ☐ | **Valid only for:**<br>Macaws (2)<br>*Anodorhynchus* spp.<br>*Ara* spp.<br>Cockatoos (2)<br>*Cacatua* spp. (2)<br>*Nymphicus hollandicus*<br>*Psittacus erithacus* | **23%** ☐ | **68%** ☐ | **24%** ☐ | **8%** ☐ |
| Sham behaviours (e.g., sham chewing, sham flying, sham bathing) | **61%** ☐ | **21%** ☐ | **Valid only for:**<br>*Agapornis* spp.<br>Cockatoos<br>*Nymphicus hollandicus*<br>*Psittacus erithacus* | **18%** ☐ | **32%** ☐ | **62%** ☐ | **6%** ☐ |
| Whole body stereotypies (head bobbing, rocking) | **71%** ☐ | **26%** ☐ | **Valid only for:**<br>Cockatoos (3)<br>*Cacatua* spp.<br>*Cacatua alba*<br>*Cacatua galerita*<br>*Cacatua ophtalmica*<br>*Calyptorhynchus banksii*<br>*Nymphicus hollandicus*<br>*Psittacus Erithacus* (2) | **3%** ☐ | **51%** ☐ | **49%** ☐ | **0%** ☐ |
| Locomotor stereotypies (route-tracing, pacing) | **87%** ☐ | **13%** ☐ | **Valid only for:**<br>*Amazona* spp.<br>*Cockatoos* (2)<br>*Cacatua* spp.<br>*Macaws* | **0%** ☐ | **51%** ☐ | **49%** ☐ | **0%** ☐ |
| **Additional indicators proposed by the experts** | Valid for all/most of the species: | Valid, but only for some species | | Not valid | Feasible for owners | Feasible, but only for experts | Not feasible |
| Hiding | ☐ | ☐ | **Valid only for:** | ☐ | ☐ | ☐ | ☐ |
| Excessive masturbation | ☐ | ☐ | **Valid only for:** | ☐ | ☐ | ☐ | ☐ |
| Tonic-clonic immobility/freezing | ☐ | ☐ | **Valid only for:** | ☐ | ☐ | ☐ | ☐ |
| Grunting | ☐ | ☐ | **Valid only for:** | ☐ | ☐ | ☐ | ☐ |
| Tremors | ☐ | ☐ | **Valid only for:** | ☐ | ☐ | ☐ | ☐ |
| Spot pecking | ☐ | ☐ | **Valid only for:** | ☐ | ☐ | ☐ | ☐ |

| | Validity | | | | Feasibility | | |
|---|---|---|---|---|---|---|---|
| **Explorative behaviours** | Valid for all/most of the species: | Valid, but only for some species | | Not valid | Feasible for owners | Feasible, but only for experts | Not feasible |
| Response to novel objects | **74%** ☐ | **18%** ☐ | **Valid only for:**<br>Neophilic/neophobic species (3)<br>*Psittacus erithacus* | **8%** ☐ | **66%** ☐ | **28%** ☐ | **6%** ☐ |
| Response to novel food items | **68%** ☐ | **16%** ☐ | **Valid only for:**<br>Australian parakeets<br>Neophilic/neophobic species (2)<br>*Psittacus erithacus*<br>South american parrots | **16%** ☐ | **68%** ☐ | **23%** ☐ | **9%** ☐ |
| Interaction with enrichment | **84%** ☐ | **11%** ☐ | **Valid only for:**<br>*Psittacus erithacus* | **5%** ☐ | **71%** ☐ | **26%** ☐ | **3%** ☐ |
| Time spent foraging | **83.3%** ☐ | **8.3%** ☐ | **Valid only for:** | **8.3%** ☐ | **59%** ☐ | **26%** ☐ | **15%** ☐ |
| Response in unfamiliar environments | **76%** ☐ | **11%** ☐ | **Valid only for:**<br>Cockatoos<br>Neophilic/neophobic species | **13%** ☐ | **49%** ☐ | **40%** ☐ | **11%** ☐ |
| **Additional indicators proposed by the experts** | Valid for all/most of the species: | Valid, but only for some species | | Not valid | Feasible for owners | Feasible, but only for experts | Not feasible |
| Response to electronic devices (TV, music...) | ☐ | ☐ | **Valid only for:** | ☐ | ☐ | ☐ | ☐ |

| | Validity | | | | Feasibility | | |
|---|---|---|---|---|---|---|---|
| **Maintenance behaviours** | Valid for all/most of the species: | Valid, but only for some species | | Not valid | Feasible for owners | Feasible, but only for experts | Not feasible |
| Amount of time spent sleeping | **91%** ☐ | **3%** ☐ | **Valid only for:** | **6%** ☐ | **90%** ☐ | **7%** ☐ | **3%** ☐ |
| Time of day (morning, afternoon, evening) spent sleeping/resting | **66%** ☐ | **6%** ☐ | **Valid only for:** | **28%** ☐ | **79%** ☐ | **14%** ☐ | **7%** ☐ |
| Posture while sleeping (e.g., on one leg, head tucked under wings) | **70%** ☐ | **9%** ☐ | **Valid only for:** | **21%** ☐ | **86%** ☐ | **10%** ☐ | **4%** ☐ |
| Daily food intake | **85%** ☐ | **6%** ☐ | **Valid only for:**<br>*Amazona* spp.<br>Cokatoos | **9%** ☐ | **80%** ☐ | **17%** ☐ | **3%** ☐ |
| Daily water consumption | **82%** ☐ | **3%** ☐ | **Valid only for:** | **15%** ☐ | **70%** ☐ | **20%** ☐ | **10%** ☐ |
| Interest in bathing | **71%** ☐ | **16%** ☐ | **Valid only for:**<br>*Amazona* spp.<br>*Anodorhynchus* spp.<br>*Ara* spp.<br>*Melopsittacus undulatus*<br>Cockatoos<br>*Cacatua* spp. | **13%** ☐ | **82%** ☐ | **18%** ☐ | **0%** ☐ |
| Preening activity (incl. time of day and frequency, duration) | **97%** ☐ | **3%** ☐ | **Valid only for:** | **0%** ☐ | **74%** ☐ | **26%** ☐ | **0%** ☐ |
| **Additional indicators proposed by the experts** | Valid for all/most of the species: | Valid, but only for some species | | Not valid | Feasible for owners | Feasible, but only for experts | Not feasible |
| Beak maintenance | ☐ | ☐ | **Valid only for:** | ☐ | ☐ | ☐ | ☐ |

| | Validity | | | | Feasibility | | |
|---|---|---|---|---|---|---|---|
| **Parrot-Human interactions** | Valid for all/most of the species: | Valid, but only for some species | | Not valid | Feasible for owners | Feasible, but only for experts | Not feasible |
| Response upon contact with caregiver | **94%** ☐ | **3%** ☐ | **Valid only for:** | **3%** ☐ | **68%** ☐ | **29%** ☐ | **3%** ☐ |
| Response upon contact with familiar person | **91%** ☐ | **3%** ☐ | **Valid only for:** | **6%** ☐ | **71%** ☐ | **23%** ☐ | **6%** ☐ |
| Response upon contact with unfamiliar person | **85%** ☐ | **3%** ☐ | **Valid only for:** | **12%** ☐ | **70%** ☐ | **30%** ☐ | **0%** ☐ |
| Facial expressions (e.g., blushing) during human interaction | **46%** ☐ | **39%** ☐ | **Valid only for:**<br>*Amazona* spp. (2)<br>Macaws (10)<br>*Ara* spp.<br>*Melopsittacus undulatus*<br>Cockatoos<br>*Nymphicus hollandicus*<br>*Probosciger aterrimus* (2)<br>*Psittacus erithacus* | **15%** ☐ | **34%** ☐ | **53%** ☐ | **13%** ☐ |
| Ruffling of feathers (e.g., nape, crown, beard) during human interaction | **64%** ☐ | **23%** ☐ | **Valid only for:**<br>*Amazona* spp. (4)<br>Cockatoos (2)<br>*Cacatua* spp.<br>*Nymphicus hollandicus*<br>Conures<br>Larger parrots<br>Macaws (2)<br>*Psittacus erithacus* (2)<br>South american parrots | **13%** ☐ | **52%** ☐ | **38%** ☐ | **10%** ☐ |
| Willingness to approach or step up on the human | **64%** ☐ | **3%** ☐ | **Valid only for:** | **33%** ☐ | **84%** ☐ | **6%** ☐ | **10%** ☐ |
| Withdrawal from human interaction | **82%** ☐ | **6%** ☐ | **Valid only for:**<br>Larger parrots | **12%** ☐ | **66.6%** ☐ | **26.6%** ☐ | **6.6%** ☐ |
| Food-related interaction (e.g., begging for food, acceptance of food from the hand, regurgitation of food to humans) | **73%** ☐ | **9%** ☐ | **Valid only for:**<br>*Amazona* spp.<br>Cockatoos<br>Conures<br>Larger parrots<br>Macaws<br>*Psittacus Erithacus* | **18%** ☐ | **65%** ☐ | **29%** ☐ | **6%** ☐ |
| Sexual related behaviours (e.g., panting, receptive posture) | **67%** ☐ | **15%** ☐ | **Valid only for:**<br>*Amazona* spp.<br>*Cacatua* spp.<br>Cockatoos<br>Larger parrots<br>Macaws<br>*Psittacus erithacus* | **18%** ☐ | **38%** ☐ | **56%** ☐ | **6%** ☐ |
| Behaviour during manual restraint (vocalizing, biting, resistance) | **55%** ☐ | **3%** ☐ | **Valid only for:** | **42%** ☐ | **45%** ☐ | **26%** ☐ | **29%** ☐ |
| Contact seeking behaviours in absence of humans (e.g., vocalization, flapping wings) | **85%** ☐ | **3%** ☐ | **Valid only for:**<br>*Amazona* spp.<br>Cockatoos | **12%** ☐ | **56%** ☐ | **38%** ☐ | **6%** ☐ |
| Preference for some humans over others | **53%** ☐ | **9%** ☐ | **Valid only for:**<br>*Amazona* spp.<br>Macaws<br>*Psittacus Erithacus* (2) | **38%** ☐ | **66%** ☐ | **9%** ☐ | **5%** ☐ |
| Initiation of contact with human being | **73.3%** ☐ | **3.3%** ☐ | **Valid only for:**<br>*Psittacus Erithacus* | **23.3%** ☐ | **70%** ☐ | **20%** ☐ | **10%** ☐ |
| Mimicry of human sounds | **39%** ☐ | **26%** ☐ | **Valid only for:**<br>*Amazona* spp.<br>*Anodorhynchus* spp.<br>*Ara* spp.<br>*Cacatua* spp.<br>*Cacatua galerita*<br>*Melopsittacus undulatus*<br>*Nymphicus hollandicus*<br>*Psittacula* spp.<br>*Psittacula krameri*<br>*Psittacus Erithacus*<br>Larger parrots | **35%** ☐ | **66.6%** ☐ | **16.6%** ☐ | **16.6%** ☐ |
| **Additional indicators proposed by the experts** | Valid for all/most of the species: | Valid, but only for some species | | Not valid | Feasible for owners | Feasible, but only for experts | Not feasible |
| Aggression towards humans linked to a specific location or perimeter | ☐ | ☐ | **Valid only for:** | ☐ | ☐ | ☐ | ☐ |

| | | | | | | | |
|---|---|---|---|---|---|---|---|
| Eating when they see the humans eating | ☐ | ☐ | **Valid only for:** | ☐ | ☐ | ☐ | ☐ |
| Expression of abnormal behaviours (e.g. toe/nail bite, feather damaging behaviours) when human is present | ☐ | ☐ | **Valid only for:** | ☐ | ☐ | ☐ | ☐ |
| Aggressive behaviours toward humans (e.g. biting, scratching, flying over) | ☐ | ☐ | **Valid only for:** | ☐ | ☐ | ☐ | ☐ |

| | **Validity** | | | | **Feasibility** | | |
|---|---|---|---|---|---|---|---|
| **Locomotor behaviours** | Valid for all/most of the species: | Valid, but only for some species | | Not valid | Feasible for owners | Feasible, but only for experts | Not feasible |
| Level of activity (time spent inactive vs active) | **88%** ☐ | **9%** ☐ | **Valid only for:** | **3%** ☐ | **68%** ☐ | **29%** ☐ | **3%** ☐ |
| Flying | **77%** ☐ | **16%** ☐ | **Valid only for:**<br>*Amazona* spp.<br>*Cacatua* spp.<br>*Calyptorhynchus banksii*<br>*Conures*<br>*Eolophus roseicapilla*<br>*Larger parrots*<br>*Lorius* spp. | **7%** ☐ | **87%** ☐ | **10%** ☐ | **3%** ☐ |
| Walking | **75%** ☐ | **16%** ☐ | **Valid only for:**<br>*Amazona* spp.<br>*Cacatua* spp.<br>*Lorius* spp. | **9%** ☐ | **83%** ☐ | **10%** ☐ | **7%** ☐ |
| Climbing | **81%** ☐ | **13%** ☐ | **Valid only for:**<br>*Amazona* spp.<br>*Cacatua* spp.<br>*Lorius* spp.<br>*Trichoglossus haematodus* | **6%** ☐ | **83%** ☐ | **10%** ☐ | **7%** ☐ |
| Hopping | **53%** ☐ | **31%** ☐ | **Valid only for:**<br>*Amazona* spp.<br>Macaws<br>*Anodorhynchus* spp.<br>Cockatoos (2)<br>*Cacatua moluccensis*<br>*Pionites* spp. (5)<br>*Psittacula* spp. | **16%** ☐ | **76%** ☐ | **7%** ☐ | **17%** ☐ |
| Time spent on the ground/bottom of the cage | **55%** ☐ | **32%** ☐ | **Valid only for:**<br>*Agapornis* spp.<br>*Anodorhynchus* spp.<br>*Melopsittacus undulatus* (2)<br>*Nymphicus hollandicus*<br>Cockatoos<br>*Psittacus erithacus* | **13%** ☐ | **81%** ☐ | **16%** ☐ | **3%** ☐ |
| Time spent in high positions (e.g., perches, grid ceiling) | **73%** ☐ | **12%** ☐ | **Valid only for:**<br>Larger parrots | **15%** ☐ | **83.3%** ☐ | **13.3%** ☐ | **3.3%** ☐ |
| Inability to fly (physical restrictions due to cage size or trimming of feathers) | **91%** ☐ | **0%** ☐ | **Valid only for:** | **9%** ☐ | **83%** ☐ | **10%** ☐ | **7%** ☐ |
| **Additional indicators proposed by the experts** | Valid for all/most of the species: | Valid, but only for some species | | Not valid | Feasible for owners | Feasible, but only for experts | Not feasible |
| Swinging | ☐ | ☐ | **Valid only for:** | ☐ | ☐ | ☐ | ☐ |

| | Validity | | | | Feasibility | | |
|---|---|---|---|---|---|---|---|
| **Social Behaviours** | Valid for all/most of the species: | Valid, but only for some species | | Not valid | Feasible for owners | Feasible, but only for experts | Not feasible |
| Aggressive behaviour toward non-mates (e.g., chasing, biting, lunging) | **75%** ☐ | **16%** ☐ | **Valid only for:** Cockatoos (2) Lorikeets | **9%** ☐ | **69%** ☐ | **31%** ☐ | **0%** ☐ |
| Aggressive behaviour toward mates (e.g., chasing, biting, lunging) | **78%** ☐ | **19%** ☐ | **Valid only for:** Cockatoos (4) *Cacatua* spp. | **3%** ☐ | **79%** ☐ | **21%** ☐ | **0%** ☐ |
| Aggressive behaviour toward chicks | **76.6%** ☐ | **6.6%** ☐ | **Valid only for:** *Amazona* spp. *Nymphicus hollandicus* Cockatoos *Psittacus erithacus* | **16.6%** ☐ | **68%** ☐ | **24%** ☐ | **8%** ☐ |
| Allopreening | **86.6%** ☐ | **6.6%** ☐ | **Valid only for:** | **6.6%** ☐ | **71%** ☐ | **29%** ☐ | **0%** ☐ |
| Time spent in vicinity of other parrots | **90%** ☐ | **3%** ☐ | **Valid only for:** *Amazona* spp. Cockatoos Macaws | **7%** ☐ | **74%** ☐ | **26%** ☐ | **0%** ☐ |
| Frequency and duration of social interaction | **91%** ☐ | **6%** ☐ | **Valid only for:** | **3%** ☐ | **59%** ☐ | **41%** ☐ | **0%** ☐ |
| Intraspecies aggression during feeding (incl. stealing of food) | **63%** ☐ | **3%** ☐ | **Valid only for:** | **34%** ☐ | **72%** ☐ | **24%** ☐ | **4%** ☐ |
| Play behaviour towards cage-mates | **97%** ☐ | **3%** ☐ | **Valid only for:** | **0%** ☐ | **69%** ☐ | **28%** ☐ | **3%** ☐ |
| Vocal communication with other birds (e.g., type and frequency) | **85%** ☐ | **9%** ☐ | **Valid only for:** | **6%** ☐ | **59%** ☐ | **34%** ☐ | **7%** ☐ |
| **Additional indicators proposed by the experts** | Valid for all/most of the species: | Valid, but only for some species | | Not valid | Feasible for owners | Feasible, but only for experts | Not feasible |
| Displacement behaviours during social interactions | ☐ | ☐ | **Valid only for:** | ☐ | ☐ | ☐ | ☐ |

| | Validity | | | | Feasibility | | |
|---|---|---|---|---|---|---|---|
| **Sexual behaviours** | Valid for all/most of the species: | Valid, but only for some species | | Not valid | Feasible for owners | Feasible, but only for experts | Not feasible |
| Courtship feeding | **78%** ☐ | **9%** ☐ | **Valid only for:** | **13%** ☐ | **79%** ☐ | **21%** ☐ | **0%** ☐ |
| Physical proximity between mates | **90%** ☐ | **10%** ☐ | **Valid only for:** | **0%** ☐ | **79%** ☐ | **21%** ☐ | **0%** ☐ |
| Mate allopreening | **85%** ☐ | **6%** ☐ | **Valid only for:** | **9%** ☐ | **83%** ☐ | **17%** ☐ | **0%** ☐ |
| Copulation | **74%** ☐ | **3%** ☐ | **Valid only for:** | **23%** ☐ | **59%** ☐ | **27%** ☐ | **14%** ☐ |
| **Additional indicators proposed by the experts** | Valid for all/most of the species: | Valid, but only for some species | | Not valid | Feasible for owners | Feasible, but only for experts | Not feasible |
| Nest building | ☐ | ☐ | **Valid only for:** | ☐ | ☐ | ☐ | ☐ |
| Nest defence | ☐ | ☐ | **Valid only for:** | ☐ | ☐ | ☐ | ☐ |
| Masturbation | ☐ | ☐ | **Valid only for:** | ☐ | ☐ | ☐ | ☐ |
| Destruction of nesting materials | ☐ | ☐ | **Valid only for:** | ☐ | ☐ | ☐ | ☐ |
| Searching for nesting areas | ☐ | ☐ | **Valid only for:** | ☐ | ☐ | ☐ | ☐ |
| Territoriality during breeding | ☐ | ☐ | **Valid only for:** | ☐ | ☐ | ☐ | ☐ |

| | Validity | | | | Feasibility | | |
|---|---|---|---|---|---|---|---|
| **Body Display** | Valid for all/most of the species: | Valid, but only for some species | | Not valid | Feasible for owners | Feasible, but only for experts | Not feasible |
| Nape and/or crown feather ruffling | **58%** ☐ | **26%** ☐ | **Valid only for:**<br>*Amazona* spp. (5)<br>Cockatoos (2)<br>Larger parrots<br>Macaws<br>*Psittacus erithacus* (2)<br>*Deroptyus* spp. (2) | **16%** ☐ | **70%** ☐ | **23%** ☐ | **7%** ☐ |
| Cheek or beard feather ruffling | **55%** ☐ | **29%** ☐ | **Valid only for:**<br>*Amazona* spp. (4)<br>*Melopsittacus undulatus*<br>Cockatoos (3)<br>*Cacatua* spp.<br>*Nymphicus hollandicus*<br>Larger parrots | **16%** ☐ | **69%** ☐ | **24%** ☐ | **7%** ☐ |
| Wing and/or leg stretch | **65%** ☐ | **0%** ☐ | **Valid only for:** | **35%** ☐ | **73%** ☐ | **10%** ☐ | **17%** ☐ |
| Scratching | **77%** ☐ | **0%** ☐ | **Valid only for:** | **23%** ☐ | **73%** ☐ | **17%** ☐ | **10%** ☐ |
| Beak grinding | **81.2%** ☐ | **9.3%** ☐ | **Valid only for:**<br>*Cacatua* spp. | **9.3%** ☐ | **84%** ☐ | **13%** ☐ | **3%** ☐ |
| Beak whipping across perch | **76%** ☐ | **3%** ☐ | **Valid only for:**<br>*Amazona* spp.<br>*Anodorhynchus* spp.<br>*Ara* spp. | **21%** ☐ | **86%** ☐ | **11%** ☐ | **3%** ☐ |
| Wing flapping | **77%** ☐ | **7%** ☐ | **Valid only for:**<br>Lorikeets<br>Lories | **16%** ☐ | **90%** ☐ | **3%** ☐ | **7%** ☐ |
| Toe/nail biting | **69%** ☐ | **12%** ☐ | **Valid only for:**<br>*Amazona* spp.<br>*Ara* spp.<br>*Psittacus erithacus* | **19%** ☐ | **80%** ☐ | **10%** ☐ | **10%** ☐ |
| Tail wagging | **68%** ☐ | **10%** ☐ | **Valid only for:**<br>*Amazona* spp.<br>Australian parakeets<br>*Psittacus erithacus* | **22%** ☐ | **83%** ☐ | **10%** ☐ | **7%** ☐ |
| Ruffling of body feathers | **77%** ☐ | **0%** ☐ | **Valid only for:** | **23%** ☐ | **87%** ☐ | **3%** ☐ | **10%** ☐ |
| Erection of crest feathers | **49%** ☐ | **36%** ☐ | **Valid only for:**<br>Cockatoos (7)<br>*Cacatua galerita*<br>*Cacatua* spp.<br>*Nymphicus hollandicus* (4)<br>*Deroptyus* spp. | **15%** ☐ | **86.6%** ☐ | **6.6%** ☐ | **6.6%** ☐ |
| **Additional indicators proposed by the experts** | Valid for all/most of the species: | Valid, but only for some species | | Not valid | Feasible for owners | Feasible, but only for experts | Not feasible |
| Beak open all the time | ☐ | ☐ | **Valid only for:** | ☐ | ☐ | ☐ | ☐ |
| Absence of any type of body display | ☐ | ☐ | **Valid only for:** | ☐ | ☐ | ☐ | ☐ |
| Pupil movements | ☐ | ☐ | **Valid only for:** | ☐ | ☐ | ☐ | ☐ |

| | Validity | | | | Feasibility | | |
|---|---|---|---|---|---|---|---|
| **Physical measurements** | Valid for all/most of the species: | Valid, but only for some species | | Not valid | Feasible for owners | Feasible, but only for experts | Not feasible |
| Body weight | **94%** ☐ | **0%** ☐ | **Valid only for:** | **6%** ☐ | **86%** ☐ | **14%** ☐ | **0%** ☐ |
| Pectoral muscle condition score | **89%** ☐ | **7%** ☐ | **Valid only for:** | **4%** ☐ | **26%** ☐ | **74%** ☐ | **0%** ☐ |
| Condition of body feathers (contour/down feathers) | **97%** ☐ | **3%** ☐ | **Valid only for:** | **0%** ☐ | **52%** ☐ | **48%** ☐ | **0%** ☐ |
| Condition of flight feathers (wing and tail) | **97%** ☐ | **0%** ☐ | **Valid only for:** | **3%** ☐ | **52%** ☐ | **45%** ☐ | **3%** ☐ |
| Beak appearance (e.g., length, shape, position) | **100%** ☐ | **0%** ☐ | **Valid only for:** | **0%** ☐ | **43%** ☐ | **57%** ☐ | **0%** ☐ |
| Cere/nare appearance (e.g. colour, shape, size) | **94%** ☐ | **3%** ☐ | **Valid only for:** *Amazona* spp. *Melopsittacus undulatus* *Nymphicus hollandicus* *Psittacus erithacus* | **3%** ☐ | **52%** ☐ | **48%** ☐ | **0%** ☐ |
| Eye appearance (e.g., half-open, discharge) | **100%** ☐ | **0%** ☐ | **Valid only for:** | **0%** ☐ | **68%** ☐ | **32%** ☐ | **0%** ☐ |
| Mentation/alertness | **94%** ☐ | **0%** ☐ | **Valid only for:** | **6%** ☐ | **68%** ☐ | **29%** ☐ | **3%** ☐ |
| Posture (e.g., fluffed appearance, weight bearing) | **97%** ☐ | **0%** ☐ | **Valid only for:** | **3%** ☐ | **52%** ☐ | **42%** ☐ | **6%** ☐ |
| Gait changes (incl. flight) | **94%** ☐ | **0%** ☐ | **Valid only for:** | **6%** ☐ | **50%** ☐ | **46%** ☐ | **6%** ☐ |
| Respiration/breathing changes (e.g., frequency, depth) | **97%** ☐ | **0%** ☐ | **Valid only for:** | **3%** ☐ | **50%** ☐ | **47%** ☐ | **3%** ☐ |
| Number of droppings | **74%** ☐ | **7%** ☐ | **Valid only for:** | **19%** ☐ | **83%** ☐ | **10%** ☐ | **7%** ☐ |
| Appearance of droppings (e.g., colour, consistency) | **94%** ☐ | **3%** ☐ | **Valid only for:** | **3%** ☐ | **67%** ☐ | **33%** ☐ | **0%** ☐ |
| **Additional indicators proposed by the experts** | Valid for all/most of the species: | Valid, but only for some species | | Not valid | Feasible for owners | Feasible, but only for experts | Not feasible |
| Respiratory effort at rest | ☐ | ☐ | **Valid only for:** | ☐ | ☐ | ☐ | ☐ |
| Respiratory effort when disturbed | ☐ | ☐ | **Valid only for:** | ☐ | ☐ | ☐ | ☐ |
| Prolapses | ☐ | ☐ | **Valid only for:** | ☐ | ☐ | ☐ | ☐ |
| Length and frequency of moulting | ☐ | ☐ | **Valid only for:** | ☐ | ☐ | ☐ | ☐ |

#### Environment-based indicators

Please remember to save your changes as you progress through the document.

For the option "Applicable, but only to some species" you can find the list of species/taxonomic groups indicated by the experts as those for which a husbandry or management condition have an influence on welfare, followed by the number of experts that indicated them (e.g. *Amazona* spp.= indicated by one expert, *Amazona* spp. (2) = indicated by two experts). For each husbandry or management condition, you can modify the list by adding or removing species/taxonomic groups. Please modify the list **only** if you select the option "Applicable, but only to some species".

**Please select** the option "Applicable, but only to some species" only if the husbandry or management condition influence the welfare:

- only in some species or taxonomic groups, regardless the context

**Please do not select** the option "Applicable, but only to some species" if the husbandry or management condition influence the welfare of all/most of the species:

- but only in specific contexts.
- even if particularly important for only some species
- except for a few of them.

Instead, please select the option "Applicable to all/most of the species"

| | Impact on welfare | | | Applicable to | | |
|---|---|---|---|---|---|---|
| **Housing** | High | Moderate | None | All/most of the species | Only some species | |
| Cage characteristics (e.g., dimension, material, bars orientation) | **97%** ☐ | **3%** ☐ | **0%** ☐ | **100%** ☐ | **0%** ☐ | **Applicable only to:** |
| Environmental temperature | **62%** ☐ | **38%** ☐ | **0%** ☐ | **88%** ☐ | **12%** ☐ | **Applicable only to:** Lowland tropical species |
| Perches' characteristics (e.g., diameter, material) | **71%** ☐ | **26%** ☐ | **3%** ☐ | **94%** ☐ | **6%** ☐ | **Applicable only to:** Heavier species |
| Position and height of the perches (e.g., in relation to feeders or human-eye level) | **62%** ☐ | **38%** ☐ | **0%** ☐ | **97%** ☐ | **3%** ☐ | **Applicable only to:** |
| Time spent out of the cage | **73%** ☐ | **21%** ☐ | **6%** ☐ | **91%** ☐ | **9%** ☐ | **Applicable only to:** |
| Position of the cage in the room | **59%** ☐ | **41%** ☐ | **0%** ☐ | **100%** ☐ | **0%** ☐ | **Applicable only to:** |
| Room where the cage is positioned (kitchen, living room, bedroom, etc.) | **68%** ☐ | **29%** ☐ | **3%** ☐ | **100%** ☐ | **0%** ☐ | **Applicable only to:** |
| Exposure to artificial light at night | **61%** ☐ | **33%** ☐ | **6%** ☐ | **94%** ☐ | **6%** ☐ | **Applicable only to:** |
| Access to outdoor spaces | **64%** ☐ | **30%** ☐ | **6%** ☐ | **94%** ☐ | **6%** ☐ | **Applicable only to:** Medium-large sized species Species that lives in the mid canopy |
| Provision of bathing opportunities | **73%** ☐ | **27%** ☐ | **0%** ☐ | **84%** ☐ | **16%** ☐ | **Applicable only to:** |

| **Additional indicators proposed by the experts** | High | Moderate | None | All/most of the species | Only some species | |
|---|---|---|---|---|---|---|
| Size of the room where the cage is positioned | ☐ | ☐ | ☐ | ☐ | ☐ | **Applicable only to:** |
| Environmental humidity | ☐ | ☐ | ☐ | ☐ | ☐ | **Applicable only to:** |
| Presence of a retreating area/room to rest, sleep or withdraw | ☐ | ☐ | ☐ | ☐ | ☐ | **Applicable only to:** |
| Presence of a platform to stand on | ☐ | ☐ | ☐ | ☐ | ☐ | **Applicable only to:** |
| Size/number of windows in the room | ☐ | ☐ | ☐ | ☐ | ☐ | **Applicable only to:** |
| Presence of a nesting area | ☐ | ☐ | ☐ | ☐ | ☐ | **Applicable only to:** |
| Frequency of cage cleaning | ☐ | ☐ | ☐ | ☐ | ☐ | **Applicable only to:** |
| Exposure to noise | ☐ | ☐ | ☐ | ☐ | ☐ | **Applicable only to:** |
| Air quality (e.g. presence of air purifier, exposure to fresh air) | ☐ | ☐ | ☐ | ☐ | ☐ | **Applicable only to:** |
| Exposure to direct sunlight/UV light | ☐ | ☐ | ☐ | ☐ | ☐ | **Applicable only to:** |

| | | | | | | |
|---|---|---|---|---|---|---|
| Artificial light characteristics (e.g. type, intensity) | ☐ | ☐ | ☐ | ☐ | ☐ | **Applicable only to:** |

| | Impact on welfare | | | Applicable to | | |
|---|---|---|---|---|---|---|
| **Parrot-Human interaction** | High | Moderate | None | All/most of the species | Only some species | |
| Time spent without presence of a human | **52%** ☐ | **45%** ☐ | **3%** ☐ | **89%** ☐ | **11%** ☐ | **Applicable only to:**<br>Cockatoos |
| Time spent on interaction with human | **50%** ☐ | **47%** ☐ | **3%** ☐ | **90%** ☐ | **10%** ☐ | **Applicable only to:**<br>Cockatoos |
| Type of interaction with human (training, mouth to beak feeding, etc.) | **74%** ☐ | **23%** ☐ | **3%** ☐ | **93%** ☐ | **7%** ☐ | **Applicable only to:**<br>*Amazona* spp.<br>*Nymphicus hollandicus*<br>Cockatoos<br>Conures<br>Macaws<br>*Psittacus erithacus* |
| Human produces loud noises and/or sudden movements | **48.4%** ☐ | **48.4%** ☐ | **3.2%** ☐ | **97%** ☐ | **3%** ☐ | **Applicable only to:**<br>Neophobic species |
| Frequency/duration of manual restraint | **88%** ☐ | **12%** ☐ | **0%** ☐ | **100%** ☐ | **0%** ☐ | **Applicable only to:** |
| Number of people in the household | **22%** ☐ | **41%** ☐ | **37%** ☐ | **93%** ☐ | **7%** ☐ | **Applicable only to:** |
| Number of people in the household that regularly interact with the parrot | **63%** ☐ | **28%** ☐ | **9%** ☐ | **97%** ☐ | **3%** ☐ | **Applicable only to:** |
| Rearing History | **79%** ☐ | **18%** ☐ | **3%** ☐ | **100%** ☐ | **0%** ☐ | **Applicable only to:** |

| | Impact on welfare | | | Applicable to | |
|---|---|---|---|---|---|
| **Nutrition** | High | Moderate | None | All/most of the species | Only some species |
| Frequency with which food is provided | **69%** ☐ | **31%** ☐ | **0%** ☐ | **100%** ☐ | **0%** ☐ **Applicable only to:** |
| Time of day (morning, afternoon, evening) at which food is provided | **36%** ☐ | **61%** ☐ | **3%** ☐ | **97%** ☐ | **3%** ☐ **Applicable only to:** |
| Variety of food items provided | **73%** ☐ | **27%** ☐ | **0%** ☐ | **100%** ☐ | **0%** ☐ **Applicable only to:** |
| Composition of the diet (quantity of fat, cholesterol, fibre, etc.) | **94%** ☐ | **3%** ☐ | **3%** ☐ | **97%** ☐ | **3%** ☐ **Applicable only to:** Cockatoos Macaws Lorikeets Lories |
| Manner/way in which food is offered to the bird (presented in a bowl, via enrichment, etc.) | **76%** ☐ | **21%** ☐ | **3%** ☐ | **100%** ☐ | **0%** ☐ **Applicable only to:** |
| Provision of supplements (multivitamin, calcium, essential fatty acids, etc.) | **44%** ☐ | **50%** ☐ | **6%** ☐ | **100%** ☐ | **0%** ☐ **Applicable only to:** |
| Frequency of cleaning the food bowls | **81%** ☐ | **16%** ☐ | **3%** ☐ | **100%** ☐ | **0%** ☐ **Applicable only to:** |
| Pellet Size | **37%** ☐ | **47%** ☐ | **16%** ☐ | **96%** ☐ | **4%** ☐ **Applicable only to:** |
| Consumption of human food | **53%** ☐ | **30%** ☐ | **17%** ☐ | **100%** ☐ | **0%** ☐ **Applicable only to:** |
| **Additional indicators proposed by the experts** | High | Moderate | None | All/most of the species | Only some species |
| Balance between provision of fresh and dried food | ☐ | ☐ | ☐ | ☐ | ☐ **Applicable only to:** |
| Location and number of feeding areas | ☐ | ☐ | ☐ | ☐ | ☐ **Applicable only to:** |
| Way in which food is stored | ☐ | ☐ | ☐ | ☐ | ☐ **Applicable only to:** |
| Availability of clean fresh water | ☐ | ☐ | ☐ | ☐ | ☐ **Applicable only to:** |
| Frequency of fresh food provision | ☐ | ☐ | ☐ | ☐ | ☐ **Applicable only to:** |

| | Impact on welfare | | | Applicable to | |
|---|---|---|---|---|---|
| **Enrichment** | High | Moderate | None | All/most of the species | Only some species |
| Provision of cognitive enrichment | **81%** ☐ | **19%** ☐ | **0%** ☐ | **97%** ☐ | **3%** ☐ **Applicable only to:**<br>*Amazona* spp.<br>Cockatoos<br>Macaws<br>*Psittacus erithacus* |
| Provision of foraging enrichment | **91%** ☐ | **9%** ☐ | **0%** ☐ | **100%** ☐ | **0%** ☐ **Applicable only to:** |
| Provision of auditory enrichments | **48%** ☐ | **48%** ☐ | **4%** ☐ | **93%** ☐ | **7%** ☐ **Applicable only to:**<br>*Psittacus erithacus* |
| Provision of visual enrichment | **56%** ☐ | **41%** ☐ | **3%** ☐ | **94%** ☐ | **6%** ☐ **Applicable only to:**<br>*Nymphicus hollandicus*<br>Cockatoos<br>*Psittacus erithacus* |
| Rotation of enrichment | **79%** ☐ | **21%** ☐ | **0%** ☐ | **100%** ☐ | **0%** ☐ **Applicable only to:** |
| Variety of enrichment provided | **79%** ☐ | **21%** ☐ | **0%** ☐ | **100%** ☐ | **0%** ☐ **Applicable only to:** |
| Amount of enrichment provided | **70%** ☐ | **30%** ☐ | **0%** ☐ | **97%** ☐ | **3%** ☐ **Applicable only to:** |
| Opportunities to do physical exercise (flying, climbing, etc.) | **91%** ☐ | **9%** ☐ | **0%** ☐ | **97%** ☐ | **3%** ☐ **Applicable only to:**<br>Parakeets |
| Provision of chewable items | **82%** ☐ | **15%** ☐ | **3%** ☐ | **87%** ☐ | **13%** ☐ **Applicable only to:**<br>*Amazona* spp.<br>Cockatoos<br>Macaws<br>*Psittacus erithacus*<br>Species that manipulate items in the wild |
| Provision of climbing toys, swings and ladders | **58%** ☐ | **36%** ☐ | **6%** ☐ | **91%** ☐ | **9%** ☐ **Applicable only to:**<br>*Melopsittacus undulatus*<br>*Nymphicus hollandicus*<br>*Forpus* spp. |
| Opportunities to select items based on preference (e.g., for colour, shape or type of material) | **75%** ☐ | **25%** ☐ | **0%** ☐ | **97%** ☐ | **3%** ☐ **Applicable only to:**<br>*Amazona* spp.<br>Cockatoos<br>Macaws<br>*Psittacus erithacus* |
| **Additional indicators proposed by the experts** | High | Moderate | None | All/most of the species | Only some species |
| Size of enrichment in relation to parrot size | ☐ | ☐ | ☐ | ☐ | ☐ **Applicable only to:** |
| Area of the room/cage where enrichment is placed | ☐ | ☐ | ☐ | ☐ | ☐ **Applicable only to:** |

| | Impact on welfare | | | | | | Applicable to | | | | |
|---|---|---|---|---|---|---|---|---|---|---|---|
| **Social needs** | High | | Moderate | | None | | All/most of the species | | Only some species | | |
| Social housing (alone vs pair vs group) | **82%** | ☐ | **15%** | ☐ | **3%** | ☐ | **97%** | ☐ | **3%** | ☐ | **Applicable only to:** |
| Opportunities for pair bonding (i.e., living with/without a mate) | **67%** | ☐ | **27%** | ☐ | **6%** | ☐ | **90%** | ☐ | **10%** | ☐ | **Applicable only to:** Pair-bonding species (2) |
| Type of social companionship (same vs different in terms of species, size, sex, origin, etc.) | **67%** | ☐ | **27%** | ☐ | **6%** | ☐ | **97%** | ☐ | **3%** | ☐ | **Applicable only to:** |
| Frequency/duration of social separation events | **82%** | ☐ | **15%** | ☐ | **3%** | ☐ | **100%** | ☐ | **0%** | ☐ | **Applicable only to:** |
| Level of social contacts (only vocal, visual and vocal, physical) | **94%** | ☐ | **6%** | ☐ | **0%** | ☐ | **100%** | ☐ | **0%** | ☐ | **Applicable only to:** |
| **Additional indicators proposed by the experts** | High | | Moderate | | None | | All/most of the species | | Only some species | | |
| Partner/cage mate choice (free vs imposed) | ☐ | | ☐ | | ☐ | | ☐ | | ☐ | | **Applicable only to:** |

#### Questions

Please remember to save your changes as you progress through the document.

In this section you are presented with a list of factors that may influence companion parrots' welfare, about which you provided your opinion in the first round. In addition, now you can see the percentage of experts that selected each option in bold (e.g. **80%**) and your previous responses indicated as ticked boxes (☒). In this second round you have the opportunity to review the general agreement between experts and your previous responses, and decide whether or not you want to alter your responses. If you decide to alter your response, please deselect the ticked box and tick the box next to your new answer.

*Mashing: process of finely grounding and mashing food so that ingredients can not be easily separated.*
*Pelleting: process of converting finely ground mash feed into dense, free flowing pellets*

| **Diet** | Answer options | | | |
|---|---|---|---|---|
| | Always Unbalanced | Always Balanced | Balanced, but only for some species | |
| A diet exclusively based on seeds is… | **81%** ☐ | **3%** ☐ | **16%** ☐ | **Balanced for:**<br>*Melopsittacus undulatus* (4)<br>*Nymphicus hollandicus* (3)<br>*Neophema* spp. |
| A diet exclusively based on one type of seed is… | **97%** ☐ | **3%** ☐ | **0%** ☐ | **Balanced for:** |
| A diet exclusively based on pellets is… | **60%** ☐ | **20%** ☐ | **20%** ☐ | **Balanced for:**<br>*Nymphicus hollandicus*<br>Medium-large sized species<br>*Psittacula krameri* |
| A diet exclusively based on mashed food is… | **67%** ☐ | **7%** ☐ | **26%** ☐ | **Balanced for:**<br>*Nymphicus hollandicus*<br>*Forpus* spp.<br>*Psittacus erithacus*<br>Species that can't eat hard food items (pellet, seeds)<br>*Brotogeris* spp. |

*In the following table, "welfare problem" refers to both behavioural and health problems. Examples of behavioural problems are excessive fear, abnormal behaviours, apathy, while examples of health problems are fungal, bacterial and viral infections, diseases and pathologic conditions.*

*Natural behaviours (e.g. loud vocalizations, chewing) that are described as problematic by the owner(s) are not included in behavioural problems.*

| **Early life and pre-acquirement experiences** | Answer options | | | | | |
|---|---|---|---|---|---|---|
| | Less likely to develop/show welfare problems | | More likely to develop/show welfare problems | | Neither more nor less likely to develop/show welfare problems | |
| Hand-reared parrots are… | **6%** | ☐ | **58%** | ☐ | **36%** | ☐ |
| Parent-reared parrots regularly human-handled after eye opening until time of weaning are… | **46.6%** | ☐ | **16.6%** | ☐ | **36.6%** | ☐ |
| Parent-reared parrots that are not human-handled in early life are… | **29%** | ☐ | **32%** | ☐ | **39%** | ☐ |
| Parrots acquired before the end of weaning are… | **14%** | ☐ | **72%** | ☐ | **14%** | ☐ |
| Parrots acquired from pet shops are… | **0%** | ☐ | **59%** | ☐ | **41%** | ☐ |
| Parrots hand-reared with siblings (vs hand-reared alone) are… | **71.4%** | ☐ | **7.4%** | ☐ | **21.4%** | ☐ |
| Wild-caught parrots sold as companion animals are… | **6%** | ☐ | **76%** | ☐ | **16%** | ☐ |
| Parrots rehomed from shelters or sanctuaries are… | **0%** | ☐ | **60%** | ☐ | **40%** | ☐ |

*Natural behaviours (e.g. loud vocalizations, chewing) that are described as problematic by the owner(s) are not included in behavioural problems.*

| **Sex/Species differences, personality, and opinion on keeping parrots as companions.** | **Answer options** | | | | | |
|---|---|---|---|---|---|---|
| | Agree | | Disagree | | Neither agree or disagree | |
| Some parrot species are more likely to develop behavioural problems when kept in captivity. | **84%** **If you select this answer, please fill Table 1** | ☐ | **3%** | ☐ | **13%** | ☐ |
| Some parrot species are more likely to develop specific diseases/pathologies when kept in captivity. | **71%** **If you select this answer, please fill Table 2** | ☐ | **6%** | ☐ | **23%** | ☐ |
| Assessing parrot personality can improve/ensure parrot welfare. | **88%** | ☐ | **0%** | ☐ | **12%** | ☐ |
| There are sex-based differences in incidence of behavioural problems in parrots. | **47%** **If you select this answer, please fill Table 3** | ☐ | **16%** | ☐ | **40%** | ☐ |
| There are sex-based differences in incidence of disease or pathology in parrots. | **61%** **If you select this answer, please fill Table 4** | ☐ | **11%** | ☐ | **28%** | ☐ |
| Some parrot species are more suitable to be kept as companion animals. | **68%** **If you select this answer, please fill Table 5** | ☐ | **12%** | ☐ | **18%** | ☐ |
| Some parrot species should not be kept as companion animals. | **79%** **If you select this answer, please fill Table 6** | ☐ | **0%** | ☐ | **21%** | ☐ |
| Parrots should not be kept as companion animals. | **33%** | ☐ | **37%** | ☐ | **30%** | ☐ |

#### Ranks

Please remember to save your changes as you progress through the document.

In this section you are presented with the list of the 10 animal-based and environment-based indicators that experts collectively indicated as the most important in the first round. Each expert individually selected and ranked the indicators most important to them, and the indicators were selected as overall most important according to their rank score, which was calculated as follows:

$$Rank\ score = \frac{i_1w_1 + i_2w_2 + i_3w_3 \ldots i_{10}w_{10}}{Total\ responses}$$

where:
$i_n$= number of experts that selected the indicator in rank position n (1, 2, 3…10)
$w_n$= weight on the rank position (e.g. n-rank position 1=10, n-rank position 10=1)

Higher scores correspond to greater considered importance.

**If you do not agree** with these ranks, please re-rank the indicators by writing their new rank position in the column "Round 2 Rank position".

**If you agree** with the current ranks, please copy and paste the numbers from column "Round 1 Rank position" to the column "Round 2 Rank position"

| Animal-based indicator | Rank score | Round 1 Rank position | Round 2 Rank position |
|---|---|---|---|
| Feather destructive behaviour (chewing, biting, fraying, plucking) | 6.00 | 1 | |
| Whole body stereotypies (head bobbing, rocking) | 3.68 | 2 | |
| Level of activity (time spent inactive vs active) | 3.52 | 3 | |
| Expression of avoidance or escape behaviours | 3.06 | 4 | |
| Locomotor stereotypies (route-tracing, pacing) | 3.06 | 4 | |
| Excessive vocalization/screaming | 2.48 | 6 | |
| Daily food intake | 2.19 | 7 | |
| Interaction with enrichment | 2.19 | 8 | |
| Inability to fly (physical restrictions due to cage size or trimming of feathers) | 1.97 | 9 | |
| Response upon contact with caregiver | 1.71 | 10 | |

| **Environment-based indicator** | **Rank score** | **Round 1 Rank position** | **Round 2 Rank position** |
|---|---|---|---|
| Time spent out of the cage | 5.23 | 1 | |
| Opportunities to do physical exercise (flying, climbing, etc.) | 4.55 | 2 | |
| Cage characteristics (e.g., dimension, material, bars orientation) | 4.45 | 3 | |
| Social housing (alone vs pair vs group) | 3.97 | 4 | |
| Provision of foraging enrichment | 3.81 | 5 | |
| Provision of cognitive enrichment | 3.10 | 6 | |
| Composition of the diet (quantity of fat, cholesterol, fibre, etc.) | 2.48 | 7 | |
| Rearing History | 2.35 | 8 | |
| Variety of enrichment provided | 2.32 | 9 | |
| Access to outdoor spaces | 2.29 | 10 | |

#### Table 1

Please fill out this table **only if** you chose the option "Agree" for the sentence "Some parrot species are more likely to develop behavioural problems when kept in captivity."

In this table you can find the species, the number of experts that indicated the species as more likely to develop behavioural problems, and the corresponding behavioural problems. Behavioural problems are followed by the number of experts that indicated them (e.g. Aggressiveness (1) = one expert indicated aggressiveness). Please provide your opinion by selecting one of the two available options (Agree, Disagree).

For example, do you agree that lovebirds (*Agapornis* spp.) are more likely to develop specific behavioural problems? In addition, do you agree that lovebirds (*Agapornis* spp.) are more likely to develop aggressiveness and feather damaging behaviour?

Please **do not tick the box** if you neither agree or disagree.

**Please keep in mind the following information while you are answering:**

-We define "cockatoos" as all the species belonging to the superfamily *Cacatuoidea*, excluding cockatiels (*Nymphicus hollandicus*). White cockatoo species (e.g. *C. sulphurea*, *C. alba*, *C. galerita*, *C. moluccensis*,) can be indicated with "*Cacatua* spp.*"*.

-We define "macaws" as all the species belonging to the genera *Ara*, *Anodorhynchus*, *Cyanopsitta*, *Orthopsittaca*, *Primolius* and *Diopsittaca*.

Please **do not tick the box** if you neither agree or disagree.

| **Species indicated as those more likely to develop behavioural problems** | **Experts that indicated the species (n)** | **Agree** | **Disagree** | **Specific behavioural problems (number of experts that mentioned them)** |
|---|---|---|---|---|
| Lovebirds (*Agapornis* spp.) | 1 | ☐ | ☐ | Aggressiveness (1): Agree ☐ Disagree ☐<br>Feather damaging behaviour (1): Agree ☐ Disagree ☐ |
| Amazon parrots (*Amazona* spp.) | 4 | ☐ | ☐ | Feather damaging behaviour (2): Agree ☐ Disagree ☐<br>Aggressiveness (2): Agree ☐ Disagree ☐<br>Unwanted vocalizations (1): Agree ☐ Disagree ☐ |
| Macaws | 4 | ☐ | ☐ | Feather damaging behaviour (3): Agree ☐ Disagree ☐<br>Excessive vocalizations (1): Agree ☐ Disagree ☐ |
| *Ara* spp. | 1 | ☐ | ☐ | Feather damaging behaviour (1): Agree ☐ Disagree ☐ |
| Red-and-green macaw (*Ara chloropterus*) | 1 | ☐ | ☐ | Aggressiveness (1): Agree ☐ Disagree ☐ |
| Scarlet macaw (*Ara macao*) | 1 | ☐ | ☐ | Aggressiveness (1): Agree ☐ Disagree ☐ |
| Military macaw (*Ara militaris*) | 1 | ☐ | ☐ | Aggressiveness (1): Agree ☐ Disagree ☐ |
| Red-fronted macaw (*Ara rubrogenys*) | 1 | ☐ | ☐ | |
| Cockatoos (excluding cockatiels) | 13 | ☐ | ☐ | Feather damaging behaviour (6): Agree ☐ Disagree ☐<br>Aggressiveness (2): Agree ☐ Disagree ☐<br>Hormonal behaviours (1): Agree ☐ Disagree ☐ |
| *Cacatua* spp. | 1 | ☐ | ☐ | Aggressiveness (1): Agree ☐ Disagree ☐<br>Feather damaging behaviour (1): Agree ☐ Disagree ☐ |
| White Cockatoo (*Cacatua alba*) | 3 | ☐ | ☐ | Feather damaging behaviour (1): Agree ☐ Disagree ☐ |

| | | | | |
|---|---|---|---|---|
| Sulphur-crested cockatoo (*Cacatua galerita*) | 2 | ☐ | ☐ | |
| Salmon-crested cockatoo (*Cacatua moluccensis*) | 2 | ☐ | ☐ | Feather damaging behaviour (1): Agree ☐ Disagree ☐ |
| Blue-eyed cockatoo (*Cacatua ophthalmica*) | 1 | ☐ | ☐ | |
| Yellow-crested cockatoo (*Cacatua sulphurea*) | 1 | ☐ | ☐ | |
| Eclectus parrot (*Eclectus roratus*) | 1 | ☐ | ☐ | Feather damaging behaviour (1): Agree ☐ Disagree ☐ |
| Golden conure (*Guaruba guarouba*) | 1 | ☐ | ☐ | Feather damaging behaviour (1): Agree ☐ Disagree ☐ |
| Monk parakeet (*Myiopsitta monachus*) | 1 | ☐ | ☐ | Feather damaging behaviour (1): Agree ☐ Disagree ☐ |
| Caiques (*Pionites* spp.) | 2 | ☐ | ☐ | Feather damaging behaviour (1): Agree ☐ Disagree ☐<br>Aggressiveness (1): Agree ☐ Disagree ☐ |
| African grey parrot (*Psittacus erithacus*) | 10 | ☐ | ☐ | Feather damaging behaviour (7): Agree ☐ Disagree ☐<br>Excessive vocalizations (1): Agree ☐ Disagree ☐<br>Phobic behaviour (1): Agree ☐ Disagree ☐ |
| Highly intelligent species | 1 | ☐ | ☐ | |
| Larger parrots | 1 | ☐ | ☐ | |
| Long-living species | 1 | ☐ | ☐ | |
| Species with long juvenile dependency | 1 | ☐ | ☐ | |

#### Table 2

In this table you can find the species, the number of experts that indicated the species as more likely to develop the diseases or pathologies, and the diseases or pathologies that a species is more likely to develop. Diseases or pathologies are followed by the number of experts that indicated them (e.g. Neoplasia (1) = one expert indicated neoplasia). Please provide your opinion by selecting one of the two available options (agree, disagree).

For example, do you agree that lovebirds (*Agapornis* spp.) are more likely to develop specific diseases or pathologies? In addition, do you agree that lovebirds (*Agapornis* spp.) are more likely to develop neoplasia and reproductive disorders?

**Please keep in mind the following information while you are answering:**

-We define "cockatoos" as all the species belonging to the superfamily *Cacatuoidea*, excluding cockatiels (*Nymphicus hollandicus*). White cockatoo species (e.g. *C. sulphurea*, *C. alba*, *C. galerita*, *C. moluccensis*,) can be indicated with "*Cacatua* spp.".

-We define "macaws" as all the species belonging to the genera *Ara*, *Anodorhynchus*, *Cyanopsitta*, *Orthopsittaca*, *Primolius* and *Diopsittaca*.

| **Species indicated as those more likely to develop diseases or pathologies** | **Experts that indicated the species (n)** | **Agree** | **Disagree** | **Specific diseases or pathologies (number of experts that mentioned them)** |
|---|---|---|---|---|
| Lovebirds (*Agapornis* spp.) | 2 | ☐ | ☐ | Neoplasia (1): Agree ☐ Disagree ☐<br>Reproductive disorders (1): Agree ☐ Disagree ☐ |
| Amazon parrots (*Amazona* spp.) | 7 | ☐ | ☐ | Atherosclerosis (1): Agree ☐ Disagree ☐<br>Obesity (5): Agree ☐ Disagree ☐<br>Cardiac diseases (1): Agree ☐ Disagree ☐<br>Internal papilloma disease (1): Agree ☐ Disagree ☐ |
| Macaws | 2 | ☐ | ☐ | Psittacine Beak and Feather disease (1): Agree ☐ Disagree ☐<br>Internal papilloma disease (1): Agree ☐ Disagree ☐ |
| Cockatoos (excluding cockatiels) | 2 | ☐ | ☐ | Psittacine Beak and Feather disease (1): Agree ☐ Disagree ☐ |
| *Cacatua* spp. | 1 | ☐ | ☐ | Sarcocystosis (1): Agree ☐ Disagree ☐ |
| Budgerigars (*Melopsittacus undulatus*) | 3 | ☐ | ☐ | Neoplasia (1): Agree ☐ Disagree ☐<br>Reproductive disorders (1): Agree ☐ Disagree ☐<br>Obesity (1): Agree ☐ Disagree ☐ |
| Cockatiels (*Nymphicus hollandicus*) | 3 | ☐ | ☐ | Reproductive disorders (1): Agree ☐ Disagree ☐<br>Egg-related problems (1): Agree ☐ Disagree ☐<br>Atherosclerosis (1): Agree ☐ Disagree ☐ |
| Kakariki (*Cyanoramphus novaezelandiae*) | 1 | ☐ | ☐ | Egg-related problems (1): Agree ☐ Disagree ☐ |

| | | | | |
|---|---|---|---|---|
| Eclectus parrot (*Eclectus roratus*) | 2 | ☐ | ☐ | Reproductive disorders (1): Agree ☐ Disagree ☐<br>Psittacine Beak and Feather disease (1): Agree ☐ Disagree ☐<br>Gastrointestinal diseases (1): Agree ☐ Disagree ☐ |
| Galah (*Eolophus roseicapilla*) | 1 | ☐ | ☐ | Obesity (1): Agree ☐ Disagree ☐ |
| Lories | 2 | ☐ | ☐ | Psittacine Beak and Feather disease (1): Agree ☐ Disagree ☐<br>Sarcocystosis (1): Agree ☐ Disagree ☐ |
| Lorikeets | 1 | ☐ | ☐ | Reproductive disorders (1): Agree ☐ Disagree ☐ |
| Monk parakeet (*Myiopsitta monachus*) | 1 | ☐ | ☐ | Atherosclerosis (1): Agree ☐ Disagree ☐ |
| African grey parrot (*Psittacus erithacus*) | 9 | ☐ | ☐ | Atherosclerosis (3): Agree ☐ Disagree ☐<br>Aspergillosis (2): Agree ☐ Disagree ☐<br>Sarcocystosis (1): Agree ☐ Disagree ☐<br>Cardiac diseases (1): Agree ☐ Disagree ☐<br>Hypocalcaemia (2): Agree ☐ Disagree ☐<br>Rhinoliths (1): Agree ☐ Disagree ☐ |
| Small species | 1 | ☐ | ☐ | Egg-related problems (1): Agree ☐ Disagree ☐ |
| Old world species | 1 | ☐ | ☐ | Atherosclerosis (1): Agree ☐ Disagree ☐ |
| Tropical species | 1 | ☐ | ☐ | Aspergillosis (1): Agree ☐ Disagree ☐ |

#### Table 3

Please fill out this table **only if** you chose the option "Agree" for the sentence "There are sex-based differences in incidence of behavioural problems in parrots."

In this table you can find the sexes, their corresponding behavioural problems as indicated by the experts in the first round and the species indicated by the experts that show the behavioural problems. Behavioural problems and species are followed by the number of experts that indicated them (e.g. Aggressiveness (1) = one expert indicated aggressiveness, *Amazona* spp. (2) = two experts indicated *Amazona*). Please provide your opinion by selecting one of the two available options (agree, disagree).

For example, do you agree that females are more likely to show hormonal aggressiveness? In addition, do you agree that females of *Eclectus roratus*, *Agapornis* spp., macaw, and/or *Psittaccula krameri* are more likely to show hormonal aggressiveness?

Please **do not tick the box** if you neither agree or disagree.

**Please keep in mind the following information while you are answering:**

-We define "cockatoos" as all the species belonging to the superfamily *Cacatuoidea*, excluding cockatiels (*Nymphicus hollandicus*). White cockatoo species (e.g. *C. sulphurea*, *C. alba*, *C. galerita*, *C. moluccensis*,) can be indicated with "*Cacatua* spp.*"*.

-We define "macaws" as all the species belonging to the genera *Ara*, *Anodorhynchus*, *Cyanopsitta*, *Orthopsittaca*, *Primolius* and *Diopsittaca*.

| Sex that is more likely to develop the behavioural problem | Behavioural problems (number of experts that mentioned them) | Mentioned species (number of experts that mentioned them) |
|---|---|---|
| Females | Aggressiveness (1): Agree ☐ Disagree ☐ | *Agapornis* spp. (1): Agree ☐ Disagree ☐ |
| | Hormonal aggressiveness (3): Agree ☐ Disagree ☐ | *Eclectus roratus* (1): Agree ☐ Disagree ☐<br>*Agapornis* spp. (1): Agree ☐ Disagree ☐<br>Macaws (1): Agree ☐ Disagree ☐<br>*Psittacula krameri* (1): Agree ☐ Disagree ☐ |
| | Hormonal behaviours (1): Agree ☐ Disagree ☐ | / |
| | Excessive eggbound (1): Agree ☐ Disagree ☐ | / |
| Males | Excessive regurgitation (1): Agree ☐ Disagree ☐ | *Eclectus roratus* (1): Agree ☐ Disagree ☐ |
| | Aggressiveness (3): Agree ☐ Disagree ☐ | *Amazona* spp. (2): Agree ☐ Disagree ☐ |
| | | Cockatoos (1): Agree ☐ Disagree ☐ |
| | Hormonal aggressiveness (2): Agree ☐ Disagree ☐ | Cockatoos (1): Agree ☐ Disagree ☐ |
| | Chronic masturbation (1): Agree ☐ Disagree ☐ | / |

#### Table 4

Please fill out this table **only if** you chose the option "Agree" for the sentence "There are sex-based differences in incidence of disease or pathology in parrots."

In this table you can find the sexes, their corresponding diseases or pathologies as indicated by the experts in the first round and the species indicated by the experts as those more likely to develop the disease or pathology. Species and disease or pathology are followed by the number of experts that indicated them (e.g. Reproductive disorders (5) = five experts indicated reproductive disorders, Lorikeets (1) = one expert indicated lorikeets). Please provide your opinion by selecting one of the available options (agree, disagree).

For example, do you agree that females are more likely to develop deranged moult? In addition, do you agree that females of *Eclectus roratus*, are more likely to develop deranged moult?

Please **do not tick the box** if you neither agree or disagree.

| Sex that is more likely to develop the disease or pathology | Diseases or pathologies (number of experts that mentioned them) | Mentioned species (number of experts that mentioned them) |
|---|---|---|
| Females | Reproductive disorders (5): Agree ☐ Disagree ☐ | Lorikeets (1): Agree ☐ Disagree ☐<br>*Agapornis* spp. (1): Agree ☐ Disagree ☐<br>*Nymphicus hollandicus* (1): Agree ☐ Disagree ☐<br>*Melopsittacus undulatus* (1): Agree ☐ Disagree ☐<br>*Pyrrhura* spp. (1): Agree ☐ Disagree ☐ |
| | Deranged moult (1): Agree ☐ Disagree ☐ | *Eclectus roratus* (1): Agree ☐ Disagree ☐ |
| | Atherosclerosis (1): Agree ☐ Disagree ☐ | / |
| | Hypercholesterolemia (1): Agree ☐ Disagree ☐ | / |
| | Cloacal prolapse (1): Agree ☐ Disagree ☐ | / |

**Table 5**

Please fill out this table **only if** you chose the option "Agree" for the sentence "Some parrot species are more suitable to be kept as companion animals."

This table contains the list of species/taxonomic groups that the experts indicated in the first round to be more suitable to be kept as companion animals, followed by the number of experts that indicated them. Please provide your opinion by selecting one of the three available options (agree, disagree, neither agree or disagree).

| Species indicated as most suitable to be kept as companion animals | Number of experts that indicated the species | Agree | Disagree | Neither agree or disagree |
|---|---|---|---|---|
| Lovebirds (*Agapornis* spp.) | 4 | ☐ | ☐ | ☐ |
| Amazon parrots (*Amazona* spp.) | 3 | ☐ | ☐ | ☐ |
| Turquoise-fronted amazon (*Amazona aestiva*) | 1 | ☐ | ☐ | ☐ |
| Orange-winged amazon (*Amazona amazonica*) | 1 | ☐ | ☐ | ☐ |
| Yellow-crowned amazon (*Amazona ochrocephala*) | 1 | ☐ | ☐ | ☐ |
| Blue-and-yellow macaw (*Ara ararauna*) | 1 | ☐ | ☐ | ☐ |
| Red-and-green macaw (*Ara chloropterus*) | 1 | ☐ | ☐ | ☐ |
| Mini macaws (*Primolius* spp.) | 1 | ☐ | ☐ | ☐ |
| Budgerigars (*Melopsittacus undulatus*) | 15 | ☐ | ☐ | ☐ |
| Conures (generic) | 5 | ☐ | ☐ | ☐ |
| *Aratinga* spp. | 2 | ☐ | ☐ | ☐ |
| *Pyrrhura* spp. | 3 | ☐ | ☐ | ☐ |
| Green-cheeked conure (*Pyrrhura molinae* ) | 1 | ☐ | ☐ | ☐ |
| White Cockatoo (*Cacatua alba*) | 1 | ☐ | ☐ | ☐ |
| Long-billed corella (*Cacatua tenuirostris*) | 1 | ☐ | ☐ | ☐ |
| Cockatiel (*Nymphicus hollandicus*) | 13 | ☐ | ☐ | ☐ |
| Parrotlets (*Forpus* spp.) | 1 | ☐ | ☐ | ☐ |
| Kakariki (*Cyanoramphus novaezelandiae*) | 3 | ☐ | ☐ | ☐ |
| Lorikeets (generic) | 1 | ☐ | ☐ | ☐ |
| Monk parakeet (*Myiopsitta monachus*) | 3 | ☐ | ☐ | ☐ |
| Caiques (*Pionites* spp.) | 2 | ☐ | ☐ | ☐ |

| | | | | |
|---|---|---|---|---|
| *Pionus* spp. | 2 | ☐ | ☐ | ☐ |
| *Poicephalus* spp. | 1 | ☐ | ☐ | ☐ |
| African grey parrot (*Psittacus erithacus*) | 2 | ☐ | ☐ | ☐ |
| Rosellas (*Platycercus* spp.) | 1 | ☐ | ☐ | ☐ |
| Ring-necked parakeet (*Psittacula krameri*) | 2 | ☐ | ☐ | ☐ |
| Medium size species | 1 | ☐ | ☐ | ☐ |
| Smaller species | 5 | ☐ | ☐ | ☐ |

#### Table 6

Please fill out this table **only if** you chose the option "Agree" for the sentence "Some parrot species should not be kept as companion animals."

In this table you can find the list of species/taxonomic groups that the experts indicated as unsuitable for keeping as companion animals, followed by the number of experts that indicated them. Please provide your opinion by selecting one of the three available options (agree, disagree, neither agree or disagree).

**Please keep in mind the following information while you are answering:**

We define "cockatoos" all the species belonging to the superfamily *Cacatuoidea*, excluding cockatiels (*Nymphicus hollandicus*). We define "macaws" the species belonging to the genera *Ara*, *Anodorhynchus*, *Cyanopsitta*, *Orthopsittaca*, *Primolius* and *Diopsittaca*.

| Species that should not be kept as companion animals | Number of experts that indicated the species | Agree | Disagree | Neither agree or disagree |
|---|---|---|---|---|
| Amazon parrots (*Amazona* spp.) | 2 | ☐ | ☐ | ☐ |
| Cockatoos | 9 | ☐ | ☐ | ☐ |
| Cockatoos (only large species) | 2 | ☐ | ☐ | ☐ |
| Eclectus parrot (*Eclectus roratus*) | 4 | ☐ | ☐ | ☐ |
| Parrotlets (*Forpus* spp.) | 1 | ☐ | ☐ | ☐ |
| Parrotlets (*Touit* spp.) | 1 | ☐ | ☐ | ☐ |
| Macaws | 6 | ☐ | ☐ | ☐ |
| Macaws (only large species) | 1 | ☐ | ☐ | ☐ |
| Lories | 1 | ☐ | ☐ | ☐ |
| Lorikeets | 1 | ☐ | ☐ | ☐ |
| Monk parakeet (*Myiopsitta monachus*) | 1 | ☐ | ☐ | ☐ |
| *Pionus* spp. | 1 | ☐ | ☐ | ☐ |
| *Poichephalus* spp. | 1 | ☐ | ☐ | ☐ |
| Red-breasted parakeet (*Psittacula alexandri* spp.) | 1 | ☐ | ☐ | ☐ |
| African grey parrot (*Psittacus erithacus* spp.) | 2 | ☐ | ☐ | ☐ |

| | | | | |
|---|---|---|---|---|
| All medium size species | 2 | ☐ | ☐ | ☐ |
| All large size species | 4 | ☐ | ☐ | ☐ |
| Critically endangered species | 2 | ☐ | ☐ | ☐ |
| Heavily trafficked species | 1 | ☐ | ☐ | ☐ |
| Rare species | 1 | ☐ | ☐ | ☐ |

#### End message

Dear Expert,

Thank you for your time in completing this survey and for participating in this project! Your responses are a valuable source of information for our research and a very much appreciated contribution to filling a gap in the scientific knowledge of parrot welfare.

Please send this file back to me without changing the filename, and do not forget to include your postal address to receive your gift!

We hope you will be willing to participate in our future expert consultation and wish you a nice day!

### Results

**Table S5.** List of the animal-based indicators that did not reach the 70% agreement for the answer options “valid for all/most of the species” and/or “feasible for owners”. Asterisk indicates animal-based indicators that were suggested by some participants in the first round of survey and therefore proposed only in the second round of the survey. Indicators accompanied with the symbol [a] reached consensus for the answer options “valid for all/most of the species” and “feasible, but only for experts”.

| Welfare dimensions | Indicator | Valid for all/most of the species % | Valid, but only for some species % | Not valid % | Feasible for owners % | Feasible, but only for experts % | Not feasible % | Tot. n. respondent for validity | Tot. n. respondent for feasibility |
|---|---|---|---|---|---|---|---|---|---|
| Abnormal and fear-related behaviours | Excessive vocalization/screaming | 70.0 | 20.0 | 10 | 66.7 | 28.6 | 4.7 | 20 | 21 |
| | Toe/nail biting | 78.9 | 5.3 | 15.8 | 65.0 | 35.0 | 0.0 | 19 | 20 |
| | Excessive chewing (e.g., wires) | 30.0 | 65.0 | 5.0 | 73.7 | 21.0 | 5.3 | 20 | 19 |
| | Sham behaviours (e.g., sham chewing, sham flying, sham bathing) | 66.6 | 16.7 | 16.7 | 33.3 | 66.7 | 0.0 | 18 | 18 |
| | Locomotor stereotypies (route-tracing, pacing) | 90.0 | 10.0 | 0.0 | 38.0 | 62.0 | 0.0 | 20 | 21 |
| | Whole body stereotypies (head bobbing, rocking) | 85.0 | 10.0 | 5.0 | 38.00 | 62.0 | 0.0 | 20 | 21 |
| | Excessive masturbation* | 73.7 | 5.3 | 21.0 | 42.1 | 47.4 | 10.5 | 19 | 19 |
| | Tonic-clonic immobility/freezing*[a] | 80.0 | 5.0 | 15.0 | 22.2 | 77.8 | 0.0 | 20 | 18 |
| | Grunting* | 22.2 | 22.2 | 55.6 | 40.0 | 20.0 | 40.0 | 18 | 15 |
| | Tremors* | 90.0 | 0.0 | 10.0 | 63.2 | 26.3 | 10.5 | 20 | 19 |
| | Spot pecking* | 88.9 | 0.0 | 11.1 | 66.7 | 27.7 | 5.5 | 18 | 18 |
| Explorative behaviours | Response to novel food items | 68.4 | 21.0 | 10.6 | 80.0 | 15.0 | 5.0 | 19 | 20 |
| | Time spent foraging | 94.4 | 5.6 | 0.0 | 63.1 | 31.6 | 5.3 | 18 | 19 |

| | | | | | | | | | |
|---|---|---|---|---|---|---|---|---|---|
| | Response in unfamiliar environments | 79.0 | 10.5 | 10.5 | 50.0 | 40.0 | 10.0 | 19 | 20 |
| | Response to electronic devices (TV, music...)* | 74.7 | 35.3 | 0.0 | 60.0 | 20 | 20 | 17 | 15 |
| Locomotor behaviours | Level of activity (time spent inactive vs active) | 94.7 | 5.3 | 0.0 | 63.2 | 36.8 | 0.0 | 19 | 19 |
| | Hopping | 52.6 | 31.6 | 15.8 | 77.8 | 5.5 | 16.7 | 19 | 18 |
| | Time spent on the ground/bottom of the cage | 55.6 | 33.3 | 11.1 | 89.5 | 10.5 | 0.0 | 18 | 19 |
| | Swinging* | 64.7 | 5.9 | 29.4 | 64.8 | 17.6 | 17.6 | 17 | 17 |
| Body displays | Nape and/or crown feather ruffling | 63.2 | 15.8 | 21.0 | 63.2 | 26.3 | 10.5 | 19 | 19 |
| | Cheek or beard feather ruffling | 64.8 | 17.6 | 17.6 | 61.1 | 27.8 | 11.1 | 17 | 18 |
| | Wing and/or leg stretch | 63.2 | 36.8 | 0.0 | 73.7 | 10.5 | 15.8 | 19 | 19 |
| | Toe/nail biting | 57.9 | 10.5 | 31.6 | 84.2 | 5.3 | 10.5 | 19 | 19 |
| | Tail wagging | 57.9 | 5.3 | 36.8 | 84.2 | 5.3 | 10.5 | 19 | 19 |
| | Ruffling of body feathers | 68.4 | 0.0 | 31.6 | 83.3 | 5.6 | 11.1 | 19 | 18 |
| | Erection of crest feathers | 35.0 | 45.0 | 20.0 | 84.2 | 5.3 | 10.5 | 20 | 19 |
| | Absence of any type of body display* | 72.2 | 0.0 | 27.8 | 17.6 | 64.7 | 17.6 | 18 | 17 |
| | Pupil movements* | 44.4 | 11.1 | 44.4 | 41.2 | 35.3 | 23.5 | 18 | 17 |
| Maintenance behaviours | Posture while sleeping (e.g., on one leg, head tucked under wings) | 63.2 | 0.0 | 36.8 | 87.5 | 6.25 | 6.25 | 19 | 16 |
| Parrot-human interactions | Facial expressions (e.g., blushing) during human interaction | 36.8 | 36.8 | 26.3 | 25.0 | 60.0 | 15.0 | 19 | 20 |
| | Ruffling of feathers (e.g., nape, crown, beard) during human interaction | 72.2 | 11.1 | 16.7 | 44.4 | 38.9 | 16.7 | 18 | 18 |
| | Willingness to approach or step up on the human | 44.4 | 5.6 | 50.0 | 83.3 | 0.0 | 16.7 | 18 | 18 |

| | | | | | | | | | |
|---|---|---|---|---|---|---|---|---|---|
| | Sexual related behaviours (e.g., panting, receptive posture) | 68.4 | 10.5 | 21.1 | 21.0 | 73.7 | 5.3 | 19 | 19 |
| | Behaviour during manual restraint (vocalizing, biting, resistance) | 52.6 | 0.0 | 47.4 | 50.0 | 16.7 | 33.3 | 19 | 18 |
| | Contact seeking behaviours in absence of humans (e.g., vocalization, flapping wings) | 94.7 | 0.0 | 5.3 | 50.0 | 45.0 | 5.0 | 19 | 20 |
| | Preference for some humans over others | 44.4 | 5.6 | 50.0 | 66.7 | 5.5 | 27.8 | 18 | 18 |
| | Initiation of contact with human being | 66.7 | 0.0 | 33.3 | 72.2 | 11.1 | 16.7 | 18 | 18 |
| | Mimicry of human sounds | 33.3 | 22.2 | 44.4 | 66.6 | 16.7 | 16.7 | 18 | 18 |
| | Aggression towards humans linked to a specific location or perimeter* | 77.8 | 0.0 | 22.2 | 61.1 | 33.3 | 5.6 | 18 | 18 |
| | Eating when they see the humans eating* | 38.9 | 0.0 | 61.1 | 85.7 | 0.0 | 14.3 | 18 | 14 |
| | Expression of abnormal behaviours (e.g. toe/nail bite, feather damaging behaviours) when human is present* | 94.7 | 5.3 | 0.0 | 57.9 | 42.1 | 0.0 | 19 | 19 |
| Social behaviours | Aggressive behaviour toward non-mates (e.g., chasing, biting, lunging) | 84.2 | 5.3 | 10.5 | 66.7 | 33.3 | 0.0 | 19 | 18 |
| | Allopreening | 89.4 | 5.3 | 5.3 | 61.1 | 38.9 | 0.0 | 19 | 18 |
| | Frequency and duration of social interaction | 94.7 | 5.3 | 0.0 | 50.0 | 50.0 | 0.0 | 19 | 18 |
| | Intraspecies aggression during feeding (incl. stealing of food) | 55.6 | 0.0 | 44.4 | 75.0 | 25.0 | 0.0 | 18 | 16 |
| | Play behaviour towards cage-mates | 100.0 | 0.0 | 0.0 | 66.7 | 33.3 | 0.0 | 19 | 18 |

| | | | | | | | | | |
|---|---|---|---|---|---|---|---|---|---|
| | Vocal communication with other birds (e.g., type and frequency) | 89.4 | 5.3 | 5.3 | 50.0 | 38.9 | 11.1 | 19 | 18 |
| | Displacement behaviours during social interactions* | 78.9 | 0.0 | 21.1 | 44.4 | 50.0 | 5.6 | 19 | 18 |
| Sexual behaviour | Copulation | 73.7 | 0.0 | 26.3 | 57.8 | 21.1 | 21.1 | 19 | 19 |
| | Nest building | 55.6 | 22.2 | 22.2 | 75.0 | 12.5 | 12.5 | 18 | 16 |
| | Masturbation | 64.7 | 0.0 | 35.3 | 50.0 | 37.5 | 12.5 | 17 | 16 |
| | Destruction of nesting materials | 47.0 | 5.9 | 47.0 | 60.0 | 13.3 | 26.7 | 17 | 15 |
| | Searching for nesting areas | 65.0 | 0.0 | 35.0 | 41.2 | 35.3 | 23.5 | 20 | 17 |
| | Territoriality during breeding | 72.2 | 0.0 | 27.8 | 64.3 | 28.6 | 7.1 | 18 | 14 |
| Body measurements | Pectoral muscle condition score[a] | 100 | 0.0 | 0.0 | 22.2 | 77.8 | 0.0 | 18 | 18 |
| | Condition of body feathers (contour/down feathers) | 100 | 0.0 | 0.0 | 47.4 | 52.6 | 0.0 | 19 | 19 |
| | Condition of flight feathers (wing and tail) | 100 | 0.0 | 0.0 | 52.6 | 47.4 | 0.0 | 19 | 19 |
| | Beak appearance (e.g., length, shape, position) | 100 | 0.0 | 0.0 | 31.6 | 68.4 | 0.0 | 19 | 19 |
| | Cere/nare appearance (e.g. colour, shape, size) | 100 | 0.0 | 0.0 | 42.1 | 57.9 | 0.0 | 19 | 19 |
| | Eye appearance (e.g., half-open, discharge) | 100 | 0.0 | 0.0 | 68.4 | 31.6 | 0.0 | 19 | 19 |
| | Mentation/alertness | 94.7 | 0.0 | 5.3 | 68.4 | 26.3 | 5.3 | 19 | 19 |
| | Posture (e.g., fluffed appearance, weight bearing) | 100 | 0.0 | 0.0 | 44.4 | 50.0 | 5.6 | 19 | 18 |
| | Gait changes (incl. flight) | 94.7 | 0.0 | 5.3 | 47.4 | 47.4 | 5.2 | 19 | 19 |
| | Respiration/breathing changes (e.g., frequency, depth) | 100 | 0.0 | 0.0 | 31.6 | 68.4 | 0.0 | 19 | 19 |
| | Appearance of droppings (e.g., colour, consistency) | 94.7 | 0.0 | 5.3 | 68.4 | 31.6 | 0.0 | 19 | 19 |
| | Respiratory effort at rest*[a] | 94.7 | 0.0 | 5.3 | 27.8 | 72.2 | 0.0 | 19 | 18 |

| | | | | | | | | | |
|---|---|---|---|---|---|---|---|---|---|
| | Respiratory effort when disturbed*[a] | 89.5 | 0.0 | 10.5 | 27.8 | 72.2 | 0.0 | 19 | 18 |
| | Prolapses* | 83.3 | 0.0 | 16.7 | 66.7 | 22.2 | 11.1 | 18 | 18 |
| | Length and frequency of moulting* | 76.5 | 0.0 | 23.5 | 35.7 | 50.0 | 14.3 | 17 | 14 |

**Table S6.** List of the environment-based indicators that did not reach the 70% agreement for both the answer options “high impact on welfare” and “applicable to all/most of the species”. Asterisk indicates environment-based indicators that were suggested by some participants in the first round of survey and therefore proposed only in the second round of the survey. Indicators accompanied with the symbol [a] reached consensus for the answer options “moderate impact on wefare” and “applicable to all/most of the species”.

| Husbandry and management condition | Indicator | High impact % | Moderate impact % | No Impact % | Applicable to all/most of the species % | Applicable only to some species % | Tot n. respondent for impact | Tot. n. respondent for applicability |
|---|---|---|---|---|---|---|---|---|
| Housing | Position of the cage in the room | 65.0 | 35.0 | 0.0 | 100 | 0.0 | 20 | 20 |
| | Exposure to artificial light at night | 52.6 | 42.1 | 5.3 | 89.5 | 10.5 | 19 | 19 |
| | Access to outdoor spaces | 63.1 | 31.6 | 5.3 | 94.7 | 5.3 | 19 | 19 |
| | Size of the room where the cage is positioned*[a] | 15.0 | 70.0 | 15.0 | 100 | 0.0 | 20 | 19 |
| | Environmental humidity* | 68.4 | 31.6 | 0.0 | 90 | 10.0 | 19 | 20 |
| | Presence of a platform to stand on* | 20.0 | 60.0 | 20.0 | 89.5 | 10.5 | 20 | 19 |
| | Artificial light characteristics (e.g. type, intensity)* | 60.0 | 35.0 | 5.0 | 100 | 0.0 | 20 | 20 |
| | Size/number of windows in the room* | 30.0 | 60.0 | 10.0 | 100 | 0.0 | 20 | 20 |
| | Presence of a nesting area* | 45.0 | 25.0 | 30.0 | 94.7 | 5.3 | 20 | 19 |
| Enrichment | Provision of auditory enrichments | 43.7 | 56.2 | 0.0 | 100 | 0.0 | 16 | 16 |
| | Provision of visual enrichment | 50.0 | 50.0 | 0.0 | 100 | 0.0 | 18 | 18 |

| | | | | | | | | |
|---|---|---|---|---|---|---|---|---|
| | Provision of climbing toys, swings, and ladders | 65.0 | 35.0 | 0.0 | 95.0 | 5.0 | 20 | 20 |
| | Size of enrichment in relation to parrot size* | 26.3 | 52.6 | 21.0 | 100 | 0.0 | 19 | 18 |
| | Area of the room/cage where enrichment is placed* | 47.4 | 36.8 | 15.8 | 100 | 0.0 | 19 | 16 |
| Parrot-human interaction | Time spent without presence of a human | 47.0 | 47.0 | 5.9 | 88.2 | 11.8 | 17 | 17 |
| | Time spent on interaction with human | 38.9 | 55.5 | 5.5 | 88.9 | 11.1 | 18 | 18 |
| | Human produces loud noises and/or sudden movements | 50.0 | 50.0 | 0.0 | 95.0 | 5.0 | 20 | 20 |
| | Number of people in the household | 5.3 | 52.6 | 42.1 | 100 | 0.0 | 19 | 18 |
| | Number of people in the household that regularly interact with the parrot | 57.9 | 31.6 | 10.5 | 100 | 0.0 | 19 | 19 |
| Nutrition | Time of day (morning, afternoon, evening) at which food is provided[a] | 25.0 | 75.0 | 0.0 | 100 | 0.0 | 20 | 20 |
| | Provision of supplements (multivitamin, calcium, essential fatty acids, etc.) | 50.0 | 45.0 | 5.0 | 100 | 0.0 | 20 | 19 |
| | Pellet Size | 42.1 | 47.4 | 10.5 | 100 | 0.0 | 19 | 17 |
| | Consumption of human food | 63.1 | 21.0 | 15.8 | 100 | 0.0 | 19 | 18 |
| | Balance between provision of fresh and dried food* | 60.0 | 35.0 | 5.0 | 100 | 0.0 | 20 | 19 |
| | Way in which food is stored* | 55.0 | 25.0 | 20.0 | 100 | 0.0 | 20 | 18 |
| Social needs | Opportunities for pair bonding (i.e., living with/without a mate) | 65.0 | 30.0 | 5.0 | 94.7 | 5.3 | 20 | 19 |

**Table S7**. List of the sentences related to diet and percentages of participants that selected the available answer options. In bold answer options that reached at least 70% of agreement.

| Sentence to complete | Always unbalanced (%) | Balanced, but only for some species (%) | Always Balanced (%) | Total number of respondents |
|---|---|---|---|---|
| A diet exclusively based on one type of seed is… | **100.0** | 0.0 | 0.0 | 19 |
| A diet exclusively based on seeds is… | **89.5** | 10.5 | 0.0 | 19 |
| A diet exclusively based on pellets is… | **70.6** | 5.9 | 23.5 | 17 |
| A diet exclusively based on mashed food is… | 68.8 | 25.0 | 6.2 | 16 |

**Table S8**. List of the sentences related to early-life and pre-acquirement experiences and percentages of participants that selected the available answer options. In bold answer options that reached at least 70% of agreement.

| Sentences to complete | More likely to develop/show welfare problems (%) | Neither more nor less likely to develop/show welfare problems (%) | Less likely to develop/show welfare problems (%) | Total number of respondents |
|---|---|---|---|---|
| Hand-reared parrots are… | **78.9** | 21.1 | 0.0 | 19 |
| Parrots hand-reared with siblings (vs hand-reared alone) are… | 0.0 | 11.8 | **88.2** | 17 |
| Parent-reared parrots regularly human-handled after eye opening until time of weaning are… | 11.8 | 35.3 | 52.9 | 17 |
| Parent-reared parrots that are not human-handled in early life are… | 38.9 | 27.8 | 33.3 | 18 |
| Parrots acquired before the end of weaning are… | **88.2** | 5.9 | 5.9 | 17 |
| Parrots acquired from pet shops are | 64.7 | 35.3 | 0.0 | 17 |
| Wild-caught parrots sold as companion animals are… | **77.8** | 16.7 | 5.6 | 18 |
| Parrots rehomed from shelters/sanctuaries are… | 64.7 | 35.3 | 0.0 | 17 |

**Table S9**. List of statements related to parrot welfare and parrot species suitability as companion animals and percentage of participants that selected the available answer options. In bold answer options that reached at least 70% of agreement.

| Statements | Agree (%) | Disagree (%) | Neither agree nor disagree (%) | Total number of respondents |
|---|---|---|---|---|
| Some parrot species are more likely to develop behavioural problems when kept in captivity | **83.3** | 5.6 | 11.1 | 18 |
| Some parrot species are more likely to develop specific diseases/pathologies when kept in captivity | 61.1 | 11.1 | 27.8 | 18 |
| Assessing parrot personality can improve/ensure parrot welfare | **100.0** | 0.0 | 0.0 | 20 |

| | | | | |
|---|---|---|---|---|
| There are sex-based differences in incidence of behavioural problems in parrots | 58.8 | 5.9 | 35.3 | 17 |
| There are sex-based differences in incidence of disease or pathology in parrots | 60.0 | 6.7 | 33.3 | 15 |
| Some parrot species are more suitable to be kept as companion animals | **75.0** | 10.0 | 15.0 | 20 |
| Some parrot species should not be kept as companion animals | **75.0** | 0.0 | 25.0 | 20 |
| Parrots should not be kept as companion animals | 30.0 | 25.0 | 45.0 | 20 |

**Table S10.** List of the 24 species or groups proposed by some participants as those more likely to develop/show behavioural problems and percentages of experts that selected the available answer options. In bold answer options that reached at least 70% of agreement.

| Species | Agree (%) | Disagree (%) | Neither agree nor disagree (%) | Total number of respondents |
|---|---|---|---|---|
| Lovebirds (*Agapornis* spp.) | 20.0 | 53.3 | 26.7 | 15 |
| Amazon parrots (*Amazona* spp.) | 66.7 | 20.0 | 13.3 | 15 |
| Macaws (generic) | 60.0 | 6.7 | 33.3 | 15 |
| *Ara* spp. | 54.4 | 13.3 | 33.3 | 15 |
| Red-and-green macaw (*Ara chloropterus*) | 33.3 | 13.3 | 53.4 | 15 |
| Scarlet macaw (*Ara macao*) | 26.6 | 26.7 | 46.7 | 13 |
| Military macaw (*Ara militaris*) | 26.6 | 26.7 | 46.7 | 13 |
| Red-fronted macaw (*Ara rubrogenys*) | 26.6 | 26.7 | 46.7 | 13 |
| Cockatoos (excluding cockatiels) | **93.3** | - | 6.7 | 15 |
| *Cacatua* spp. | 53.4 | 13.3 | 33.3 | 15 |
| White Cockatoo (*Cacatua alba*) | 40 | 6.7 | 53.3 | 15 |
| Sulphur-crested cockatoo (*Cacatua galerita*) | 40 | 6.7 | 53.3 | 15 |
| Salmon-crested cockatoo (*Cacatua moluccensis*) | 46.7 | 0.0 | 53.3 | 15 |
| Blue-eyed cockatoo (*Cacatua ophthalmica*) | 40.0 | 6.7 | 53.3 | 15 |
| Yellow-crested cockatoo (*Cacatua sulphurea*) | 46.7 | 0.0 | 53.3 | 15 |
| Eclectus parrot (*Eclectus roratus*) | 20.0 | 20.0 | 60.0 | 15 |
| Golden conure (*Guaruba guarouba*) | 14.3 | 35.7 | 50.0 | 14 |
| Monk parakeet (*Myiopsitta monachus*) | 13.3 | 40.0 | 46.7 | 15 |

| | | | | |
|---|---|---|---|---|
| Caiques (*Pionites* spp.) | 20.0 | 33.3 | 46.6 | 15 |
| African grey parrot (Psittacus erithacus) | **86.7** | 0.0 | 13.3 | 15 |
| Highly intelligent species | 46.7 | 6.6 | 46.7 | 15 |
| Larger parrots | 33.3 | 20.0 | 46.7 | 15 |
| Long-living species | 33.3 | 20 | 46.7 | 15 |
| Species with long juvenile dependency | 33.3 | 20 | 46.7 | 15 |

**Table S11**. Species proposed by some participants as those more likely to develop or show behavioural problems, corresponding behavioural problems proposed by participants, and percentage of participants that selected the available answer options. In bold answer options that reached at least 70% of agreement.

| **Species** | **Behavioural problem** | **Agree (%)** | **Disagree (%)** | **Neither agree nor disagree (%)** | **Total number of respondents** |
|---|---|---|---|---|---|
| Cockatoos (excluding cockatiels) | Feather damaging behaviours | **100** | 0.0 | 0.0 | 14 |
| | Aggressiveness | **92.8** | 7.2 | 0.0 | 14 |
| | Hormonal behaviours | **100** | 0.0 | 0.0 | 14 |
| African grey parrot (*Psittacus erithacus*) | Feather damaging behaviours | **100** | 0.0 | 0.0 | 13 |
| | Excessive vocalizations | 38.4 | 23.1 | 38.4 | 13 |
| | Phobic behaviours | 61.5 | 23.1 | 15.4 | 13 |

**Table S12**. List of the 28 species or groups proposed by some participants during the first round of survey as the most suitable to be kept as companion animals and percentages of participants who selected the available answer options during the second round of survey. In bold answer options that reached at least 70% of agreement.

| **Species** | **Agree (%)** | **Disagree (%)** | **Neither agree nor disagree (%)** | **Total number of respondents** |
|---|---|---|---|---|
| Lovebirds (*Agapornis* spp.) | **86.7** | 6.7 | 6.6 | 15 |
| Amazon parrots (*Amazona* spp.) | 15.4 | 53.8 | 30.8 | 13 |
| Turquoise-fronted amazon (*Amazona aestiva*) | 23.1 | 38.5 | 38.4 | 13 |
| Orange-winged amazon (*Amazona amazonica*) | 23.1 | 38.5 | 38.4 | 13 |
| Yellow-crowned amazon (*Amazona ochrocephala*) | 23.1 | 38.5 | 38.4 | 13 |
| Blue-and-yellow macaw (*Ara ararauna*) | 15.4 | 61.5 | 23.1 | 13 |
| Red-and-green macaw (*Ara chloropterus*) | 0.0 | 69.2 | 30.8 | 13 |
| Mini macaws (*Primolius* spp.) | 23.1 | 38.5 | 38.4 | 13 |
| Budgerigars | **100** | 0.0 | 0.0 | 15 |

| | | | | |
|---|---|---|---|---|
| (*Melopsittacus undulatus*) | | | | |
| Conures (generic) | 57.1 | 14.3 | 28.6 | 14 |
| *Aratinga* spp. | 46.1 | 15.4 | 38.5 | 13 |
| *Pyrrhura* spp. | **76.9** | 0.0 | 23.1 | 13 |
| Green-cheeked conure (*Pyrrhura molinae*) | **76.9** | 0.0 | 23.1 | 13 |
| White Cockatoo (*Cacatua alba*) | 0.0 | **92.9** | 7.1 | 14 |
| Long-billed corella (*Cacatua tenuirostris*) | 0.0 | **78.6** | 21.4 | 14 |
| Cockatiel (*Nymphicus hollandicus*) | **93.3** | 6.7 | 0.0 | 15 |
| Parrotlets (*Forpus* spp.) | **78.6** | 0.0 | 21.4 | 14 |
| Kakariki (*Cyanoramphus novaezelandiae*) | 53.8 | 7.7 | 38.5 | 13 |
| Lorikeets (generic) | 7.7 | 30.8 | 61.5 | 13 |
| Monk parakeet (*Myiopsitta monachus*) | **76.9** | 15.4 | 7.7 | 13 |
| Caiques (*Pionites* spp.) | 57.1 | 14.3 | 28.6 | 14 |
| *Pionus* spp. | 28.6 | 21.4 | 50.0 | 14 |
| *Poicephalus* spp. | 21.4 | 28.6 | 50.0 | 14 |
| African grey parrot (*Psittacus erithacus*) | 14.3 | 64.3 | 21.4 | 14 |
| Rosellas (*Platycercus* spp.) | 42.8 | 28.6 | 28.6 | 14 |
| Ring-necked parakeet (*Psittacula krameri*) | 64.3 | 7.1 | 28.6 | 14 |
| Medium size species | 15.4 | 7.7 | **76.9** | 13 |
| Smaller species | 66.7 | 6.6 | 26.7 | 15 |

**Table S13.** List of the 20 species or groups proposed by some participants during the first round of survey as those that should not be kept as companion animals and percentages of participants who selected the available answer options during the second round of survey. In bold answer options that reached at least 70% of agreement.

| Species | Agree (%) | Disagree (%) | Neither agree nor disagree (%) | Total number of respondents |
|---|---|---|---|---|
| Amazon parrots (*Amazona* spp.) | 53.4 | 13.3 | 33.3 | 15 |
| Cockatoos (excluding cockatiels) | **86.7** | 0.0 | 13.3 | 15 |
| Cockatoos (only large species) | **85.7** | 0.0 | 14.3 | 14 |
| Eclectus parrot (*Eclectus roratus*) | 62.5 | 12.5 | 25.0 | 16 |
| Parrotlets (Forpus spp.) | 6.6 | 66.7 | 26.7 | 15 |
| Parrotlets (*Touit* spp.) | 20.0 | 40.0 | 40.0 | 15 |
| Macaws | 68.8 | 18.8 | 12.4 | 16 |
| Macaws (only large species) | 50.0 | 28.6 | 21.4 | 14 |
| Lories | 53.4 | 13.3 | 33.3 | 15 |
| Lorikeets | 46.7 | 13.3 | 40.0 | 15 |
| Monk parakeet (*Myiopsitta monachus*) | 21.4 | 42.9 | 35.7 | 14 |

| | | | | |
|---|---|---|---|---|
| *Pionus* spp. | 26.7 | 13.3 | 60.0 | 15 |
| *Poichephalus* spp. | 26.7 | 26.6 | 46.7 | 15 |
| Red-breasted parakeet (*Psittacula alexandri*) | 13.3 | 33.3 | 53.4 | 15 |
| African grey parrot (*Psittacus erithacus*) | 56.3 | 12.4 | 31.3 | 16 |
| All medium size species | 13.3 | 40.0 | 46.7 | 15 |
| All large size species | 43.8 | 18.7 | 37.5 | 16 |
| Critically endangered species | **75.0** | 6.2 | 18.8 | 16 |
| Heavily trafficked species | **81.3** | 6.2 | 12.5 | 16 |
| Rare species | 68.8 | 6.2 | 25.0 | 16 |
